# Enhanced kinase translocation reporters for simultaneous real-time measurement of PKA, ERK, and Ca^2+^

**DOI:** 10.1101/2024.09.30.615856

**Authors:** Shang-Jui Tsai, Yijing Gong, Austin Dabbs, Fiddia Zahra, Junhao Xu, Aleksander Geske, Michael J. Caterina, Stephen J. Gould

**Affiliations:** Department of Neurosurgery, Johns Hopkins University School of Medicine, Baltimore, MD 21205 USA; Department of Neuroscience, Johns Hopkins University School of Medicine, Baltimore, MD 21205; Department of Biological Chemistry, Johns Hopkins University School of Medicine, Baltimore, MD 21205 USA

## Abstract

Kinase translocation reporters (KTRs) are powerful tools for single-cell measurement of time-integrated kinase activity but suffer from restricted dynamic range and limited sensitivity, particularly in neurons. To address these limitations, we developed enhanced KTRs (eKTRs) for protein kinase A (PKA) and extracellular signal-regulated kinase (ERK) that display high sensitivity, rapid response kinetics, broad dynamic range, cell type-specific tuning, and an ability to detect PKA and ERK activity in primary sensory neurons. Moreover, co-expression of optically separable eKTRs for PKA and ERK revealed the kinetics of expected and unexpected crosstalk between PKA, ERK, protein kinase C, and calcium signaling pathways, demonstrating the utility of eKTRs for live cell monitoring of diverse and interacting signaling pathways. These results open the door to improved live-cell and in vivo measurements of key signaling pathways in neurons, while at the same time demonstrating the importance of KTR size and NLS strength to KTR dynamics.

## INTRODUCTION

Protein kinase signaling pathways play critical roles in biology. In eukaryotes, protein kinase A (**PKA**) and extracellular signal-regulated kinase (**ERK**) are particularly important, as PKA is a key effector of the second messenger cyclic AMP (**cAMP**)(*1–4*), while growth factor signaling, exemplified by epidermal growth factor (**EGF**) activation of EGF receptors (**EGFR**), activates ERK (*5, 6*). A variety of fluorescent protein-based kinase biosensors have been developed, to monitor kinase activities in living cells, and these have yielded important new insights into the PKA and ERK signaling pathways in individual living cells(*7–11*). These biosensors range from non-integrative sensors that detect changes in kinase activity on the seconds-to-minutes timescale, to transcription-based sensors that report kinase activities on the timescale of hours-to-days (*12–23*). Between these two extremes, the kinase translocation reporters (**KTRs**) provide an integrative measurement of kinase activation and inhibition on the critical minutes-to-hour timescale that corresponds to many physiological processes(*24, 25*).

As originally conceived(*24*), KTRs are comprised of a fluorescent protein fused to a kinase sensor domain with four functional components: (***i***) a kinase binding site, (***ii***) a bipartite nuclear localization signal (**bNLS**), (***iii***) a nuclear export signal (**NES**), and (***iv***) a pair of kinase phosphorylation sites, one in the central region of the bNLS and the other positioned between the bNLS and NES(*24*). At rest, the activity of the NLS dominates over that of the NES, and the KTR is localized primarily to the nucleus. In contrast, kinase activation leads to KTR phosphorylation, with the increased negative charge inhibiting NLS activity, leading to nuclear export of the KTR and a sharp rise in the cytoplasm/nucleus (C/N) ratio of KTR fluorescence that’s easily detected by widefield microscopy.

Given that KTRs are integrative biosensors that operate at biologically meaningful timescales, their optimization is of high priority. This is particularly true for PKA KTRs, which display relatively narrow dynamic ranges and poor sensitivity, leaving a gap in our ability to follow minutes-to-hours integrative changes in PKA activity in real time. Using PKA-KTR as a proof-of-principle example, we show that KTR performance characteristics can be improved significantly by (1) increasing KTR size above the ∼40 kDa diffusion limit of the nuclear pore complex, and (2) increasing NLS strength, both of which ensure that KTR localization is determined by a continual competition between active nuclear import and active export. Furthermore, we show that the enhanced KTRs described in this report are able to monitor PKA or ERK in neurons whereas previously reported KTRs for PKA and ERK were not. Furthermore, we show that multiplexing eKTRs for PKA and ERK together with GCaMP8 allows for the simultaneous measurement of PKA, ERK and calcium in live cells using nothing more sophisticated than widefield fluorescence microscopy

## RESULTS

### Tuning bNLS strength to improve the sensitivity and dynamic range of PKA-KTR1

As noted above, the kinase sensor domain of a KTR should contain an NLS that is strong enough to drive its active nuclear protein import across the barrier of the nuclear pore complex. Given that previously described KTRs are ∼30-35 kDa, well below the ∼40 kDa diffusion size limit of the nuclear pore(*26, 27*), we reasoned that diffusion-mediated nuclear entry of KTRs might have confounded earlier efforts at KTR engineering and optimization. In particular, the issue of KTR size might explain why the original PKA-KTR(*24*), which we refer to as PKA-KTR1, displays relatively poor sensitivity to changes in PKA aactivity(*28, 29*). To explore this possibility, we appended the kinase sensor domain of PKA-KTR1 to the N-terminus of tdCherry, a tandem dimer of mCherry, creating a PKA-KTR1/tdCherry fusion protein of ∼64 kDa, which is larger than the rapid diffusion size limit of the nuclear pore(*26*). When expressed in HEK293 cells, PKA-KTR1/tdCherry was localized almost exclusively in the cytoplasm (C/N ratio of ∼1.7), even after we added forskolin (**Fsk,** 30 μM), which activates adenylate cyclase and activates PKA(*1–4*) (***Fig. S1A, B***). These results indicate that the NLS in the PKA-KTR1 sensor domain was simply too weak to drive protein import into the nucleus. Interestingly, when we expressed the a ∼34 kDa PKA-KTR1/mCherry protein, we found that it had a C/N ratio of ∼1 (***Fig. S1C, D***), consistent with the idea that diffusion plays a major role in the subcellular distribution of KTRs that are smaller than the rapid diffusion limit of the nuclear pore.

Our finding that the bNLS in PKA-KTR1 was too weak to drive nuclear import of PKA-KTR1/tdCherry led us to interrogate the strength of this bNLS sequence using the cNLS Mapper algorithm, which scored the strength of the bNLS in the PKA-KTR1 sensor domain as <5 (***Fig. 1A***). To determine whether increasing NLS strength might convert PKA-KTR1 into a good PKA biosensor, we used cNLS Mapper to design a series of PKA sensor domains of increasing NLS strength. When these domains were appended to the N-terminus of tdCherry and expressed in HEK293 cells, we found that their subcellular distributions matched their predicted NLS strengths. Specifically, ePKA-KTR1.1, which had a relatively weak cNLS Mapper score of 8 was predominantly cytoplasmic, whereas ePKA-KTRs 1.2-1.5, which had stronger cNLS Mapper scores of 10.5-15.5, all displayed C/N ratios <1 (***Fig. 1B, C***). Not surprisingly, addition of Fsk failed to increase the C/N ratio of either ePKA-KTR1.1 or ePKA-KTR1.5, which lie at the opposing ends of the ‘NLS too weak’ and ‘NLS too strong’ spectrum of ePKA-KTRs (***Fig. 1B, 1C***). In contrast, ePKA-KTR1.2, 1.3, and 1.4 all displayed a significant Fsk-induced increases in their C/N ratio. Importantly, we also found that leptomycin B (LMB), an inhibitor of nuclear protein export (*30, 31*), triggered a drop in the C/N ratios of ePKA-KTR1.2, 1.3, and 1.4 (***Fig. 1D***), indicating that their steady-state distribution involved active nuclear export, in addition to active nuclear import.

**Figure 1.**
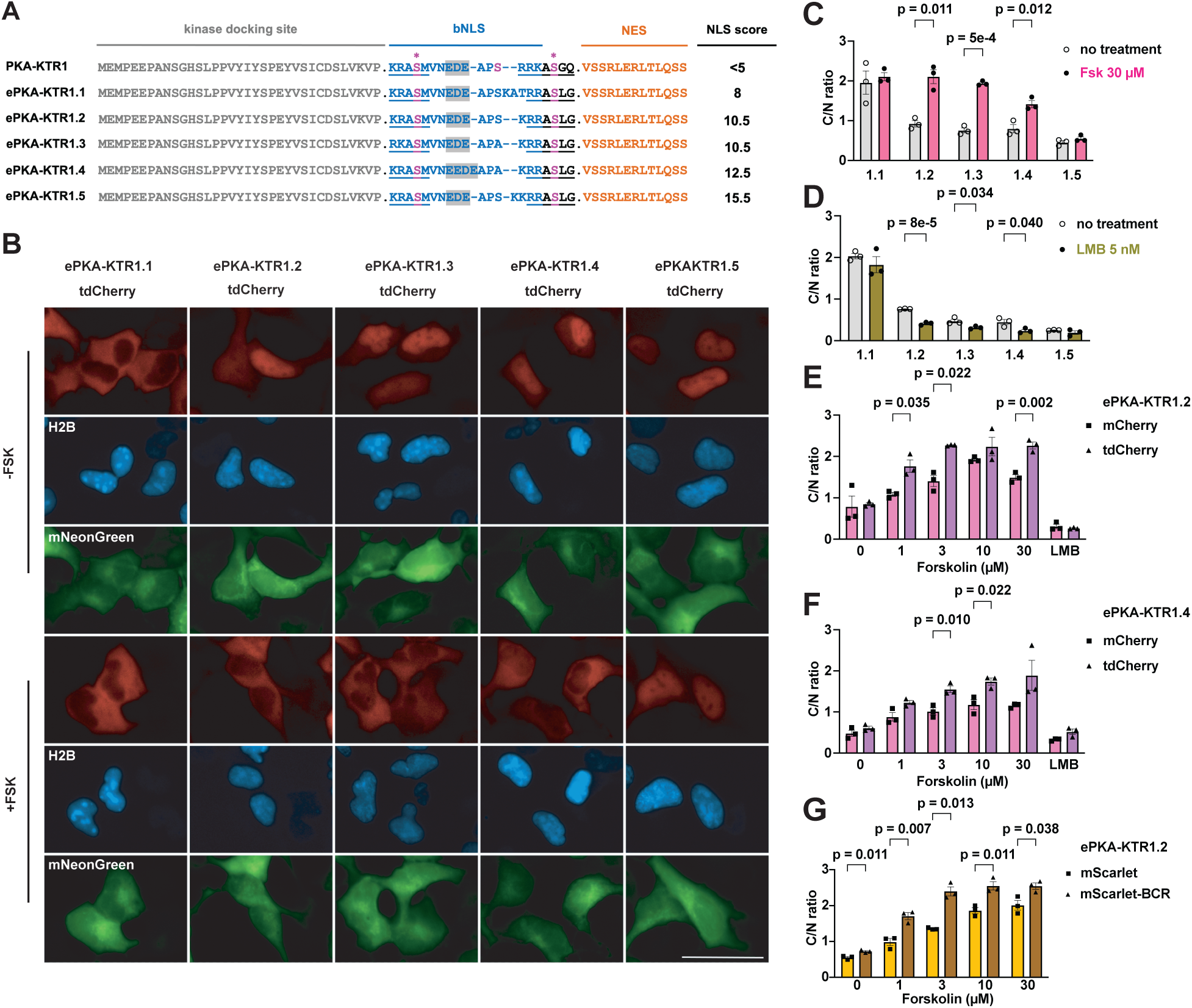
Tuning NLS strength and KTR size improves PKA-KTR sensitivity and dynamic range. (**A**) Amino acid sequences of PKA-KTR sensor domains, showing the kinase docking site, bNLS, PKA phosphorylation sites (in pink, with asterisk), and NES, with predicted NLS strength scores listed to the right. Bar, 50 μm. (**B**) Representative fluorescence microscopy images of HEK293 cells expressing the (red) ePKA-KTR1.1, 1.2, 1.3, 1.4, and 1.5 proteins, (blue) H2B-mTagBFP2, and (green) mNeonGreen, grown in the absence of Fsk (-FSK) or after 30 minutes exposure to 30 µM forskolin (+FSK). (**C**) Bar graph showing C/N ratios of ePKA-KTR1.1, 1.2, 1.3, 1.4, and 1.5/tdCherryproteins (grey) prior to the addition of Fsk or (pink) 30 minutes after the addition of Fsk. (**D**) C/N ratios of ePKA-KTR1.1, 1.2, 1.3, 1.4, and 1.5/tdCherry (grey) prior to the addition of leptomycin B or (green) after 30 minutes incubation in leptomycin B. (**E**) Bar graphs comparing the C/N ratios of ePKA-KTR1.2/mCherry (pink) and ePKA-KTR1.2/tdCherry (purple) following 30 minutes incubation in 0, 1, 3, 10, or 30 µM Fsk, or 5 nM leptomycin B (LMB) (**F**) Bar graphs comparing the C/N ratios of (pink) ePKA-KTR1.4/mCherry and (purple) ePKA-KTR1.4/tdCherry following 30 minutes incubation in 0, 1, 3, 10, or 30 µM Fsk, or 5 nM leptomycin B (LMB). (**G**) Bar graphs comparing the C/N ratios of ePKA-KTR1.2/mScarlet (gold) or ePKA-KTR1.2/mScarlet/BCR (brown) following 30 minutes incubation in 0, 1, 3, 10, or 30 µM Fsk, or 5 nM leptomycin B (LMB). Data are from a minimum of three independent biological replicates. *p* values were calculated by two-way ANOVA.

### KTR size impacts NLS-optimized KTR dynamic range

Given that protein size was critical to the design and testing of ePKA-KTRs, we next tested whether it might also affect the way that well-designed eKTR sensor domains actually performed. To explore this idea, we compared their Fsk responsiveness of eKTRs comprised of the ePKA-KTR1.2 and ePKA-KTR1.4 sensor domains to the N-terminus of either mCherry (∼34 kDa), or tdCherry (∼64 kDa) proteins. These four sensors were expressed in HEK293 cells, the cells were imaged by fluorescence microscopy before and after Fsk addition, and the resting and Fsk-induced C/N ratios were calculated. The Fsk-induced C/N ratios of the ∼64 kDa ePKA-KTRs were, in general, higher than those of the Fsk-induced C/N ratios than the small, ∼34 kDa proteins (***Fig. 1E, 1F***), consistent with idea that small KTRs diffuse from the cytoplasm into the nucleus even after phosphorylation, reducing their C/N dynamic range. These results also showed that the ePKA-KTR1.2/tdCherry sensor (NLS score = 10.5) responded well to low Fsk (1 µM) but reached saturation above 3 µM Fsk, whereas ePKA-KTR1.4/tdCherry (NLS score = 12.5) displayed a muted response to low Fsk (1 µM) but continued to increase its C/N ratio at higher Fsk concentrations (***Fig 1E, 1F***). Thus, it may be that eKTRs of varying NLS strength report at different ranges of kinase activity, with ‘weaker NLS’ eKTRs more sensitive to lower ranges of kinase activity and ‘stronger NLS’ eKTRs more sensitive at higher range of kinase activity.

### Effect of oligomerization on the behavior of a small KTR

The effect of KTR size on KTR performance characteristics led us to next test whether similar effects might be achieved by appending small oligomerizing peptides to the C-terminus of small KTRs. Towards this end, we created eKTRs comprised of the ePKA-KTR1.2 sensor domain fused to the N-terminus of mScarlet-I (*32*) and mScarlet-I/BCR, which carries the BCR homo-oligomerization peptide at its C-terminus (the BCR peptide forms anti-parallel homodimers and homotetramers(*33*)). These proteins are expected to have native size of ∼36 kDa vs ∼70-140 kDa, respectively, and when we expressed them in HEK293 cells and measured their C/N ratios, we found that the C/N ratio of ePKA-KTR1.2/mScarlet-I/BCR was higher than that of ePKA-KTR1.2/mScarlet-I, especially at the lower range of Fsk concentrations (1 or 3 µM) (***Fig. 1G***). It should be noted that native ePKA-KTR1.2/mScarlet-I/BCR dimers and tetramers will possess two or four copies of its kinase sensor domain, and thus, their C/N ratios are likely to reflect differences in both size and sensor domain copy number.

### Increasing NLS strength and oligomerization enhances ERK-KTR dynamic range

We next tested whether size and NLS strength might also improve on the ERK-KTR(*24*), which we refer to here as ERK-KTR1. ERK-KTR1 has proven to be a valuable sensor of ERK activity in multiple studies(*8, 34, 35*), has a cNLS Mapper(*36*) score of 12.5 (***Fig. 2A***), and when appended to the N-terminus of tdCherry operated as expected, specifying nuclear localization at baseline and moving to the cytoplasm in response to EGF, which activates the ERK pathway (***Fig. 2B, C***). However, cell type-specific differences in ERK kinase activity and protein import/export dynamics (*37, 38*) suggest that researchers might need eERK-KTRs tuned to different sensitivities, and we therefore built a range of eERK-KTRs with NLSs of various strengths. When expressed in HEK293 cells, the new eERK-KTR1.1, 1.2, and 1.3 proteins all displayed low C/N ratios at baseline and translocated to the cytoplasm in response to EGF (***Fig. 2B, C***), though eERK-KTR1.2 appeared to be particularly responsive to low EGF concentration. Sensitivity to low level EGF concentrations was even more notable for eERK-KTR1.2/mScarlet-I/BCR (***Fig. 2D***), perhaps because it’s oligomeric configuration would contain 2-4 copies of the eERK1.2 sensor domain.

**Figure 2.**
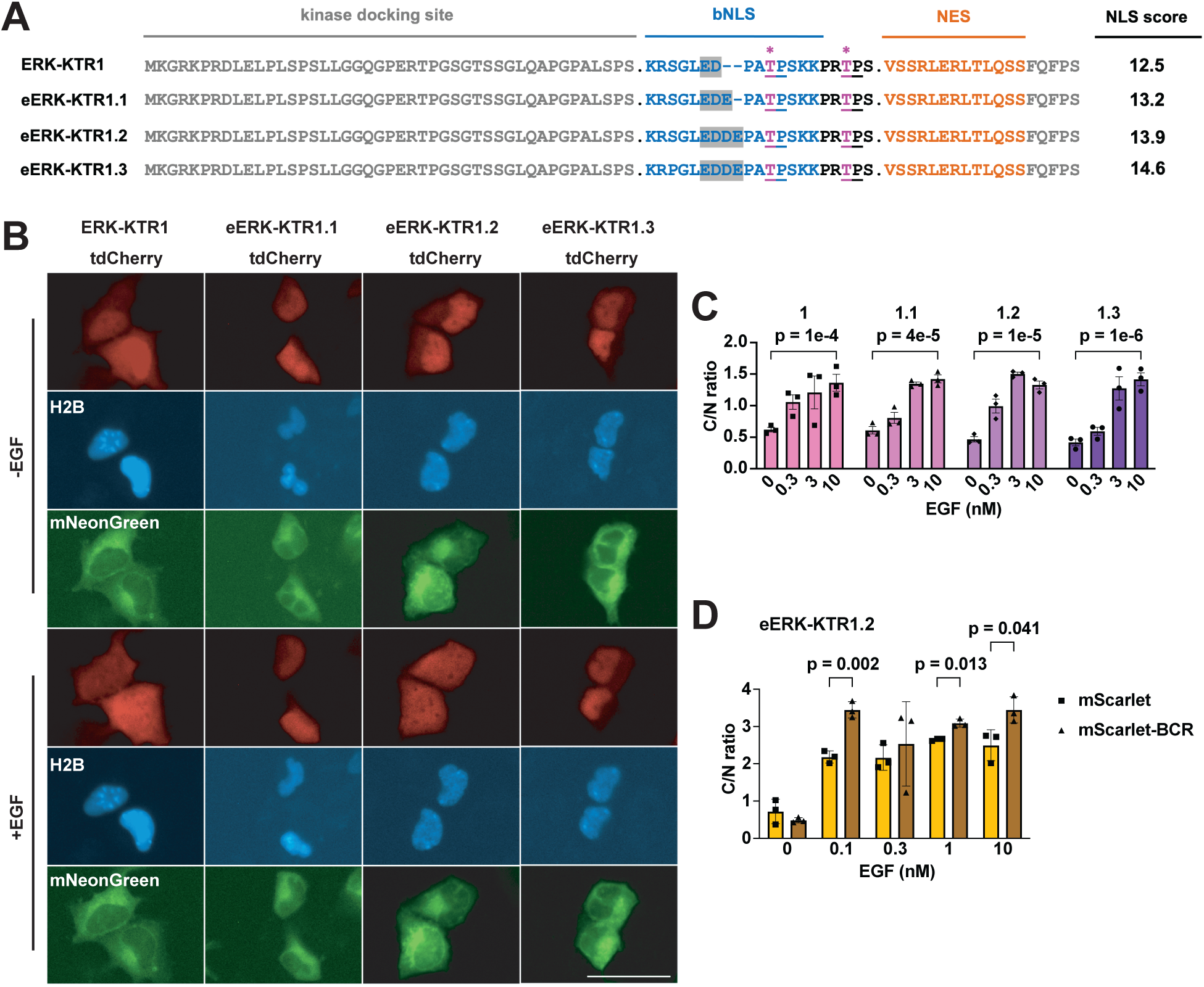
Tuning the NLS of ERK-KTR. (**A**) Amino acid sequence alignments of ERK-KTR sensor domains, showing the kinase docking site, bNLS, (pink, asterisk) PKA phosphorylation sites, and NES, with predicted NLS strength scores listed to the right. Bar, 50 μm. (**B**) Representative live-cell fluorescence images of HEK293 cells expressing H2B-mTagBFP2, mNeonGreen, and variants of ERK-KTR/tdCherry proteins. -EGF refers to fluorescence images taken prior to the addition of EGF, while +EGF refers to fluorescence micrographs taken 15 minutes after the addition of 10 nM EGF. (**C**) Bar graph showing C/N ratios of eERK-KTR1.1, 1.2, and 1.3 /tdCherry1, following 15 minutes incubation in 0, 0.3, 3, or 10 nM EGF. (**D**) C/N ratios of eERK-KTR1.2/mScarlet (gold) or eERK-KTR1.2/mScarlet/BCR (brown), after 15 minutes incubation in 0, 0.1, 0.3, 1, or 10 uM Fsk. Data are from a minimum of three independent biological replicates. *p* values were calculated by two-way ANOVA.

### Marking cytoplasm and nucleus with ER-mTagBFP2

In the course of our experiments, we found that the H2B-mTagBFP2 fusion protein that we initially used to mark the nucleus had an unexpected effect on cell size. To avoid these effects, we generated an endoplasmic reticulum (**ER**)-localized form of mTagBFP2 that carries a secretory signal sequence(*39, 40*) at its N-terminus of mTagBFP2 and a KDEL ER retrieval signal at its C-terminus(*41*). When this ER-mTagBFP2 marker was expressed in HEK293 cells, it was localized to both the nuclear envelope and to ER tubules that extend throughout the cytoplasm, allowing us to definitively mark both the nucleus and the cytoplasm with just one fluorescent protein (***fig. S2A***). Furthermore, when we measured the C/N ratio of ePKA-KTR1.2/tdCherry in cells co-expressing this and other nuclear marker proteins, H2B-mTagBFP2 or 3xNLS-mTagBFP2 markers, we found that the calculated C/N ratio of ePKA-KTR1.2/tdCherry in cells expressing ER-mTagBFP2 were slightly higher than in cells expressing either H2B-mTagBFP2 or 3xNLS-mTagBFP2 (***fig. S2B***), raising the possibility that NLS-containing nuclear markers might interfere with the nuclear protein import/export pathways on which KTRs depend.

### Kinetics, specificity, and sensitivity of ePKA-KTR1.2

We next measured the responsiveness of eKTR-KTR1.2 to known ERK activators and inhibitors. Using an HEK293 cell line that stably expresses ePKA-KTR1.2/tdTomato and ER-mTagBFP2, we confirmed that the ePKA-KTR1.2/tTomato reporter was properly localized to the nucleus, displayed a C/N ratio of ∼0.6, and responded to Fsk stimulation (3 µM) by translocating to the cytoplasm (***Fig. 3A***). This response was rapid, reaching significance within 4 minutes (1.5-fold increase, *p* = 0.04), half-maximum at 8 minutes (2.7-fold increase, *p* = 0.04), and saturation at 20 minutes (4-fold increase, *p* = 0.0005), after which it remained relatively constant (***Fig. 3B***). When cells were stimulated with Fsk (3 µM) at t = 3 min, and subsequently exposed to the PKA inhibitor H89 (50 µM) at t = 23 min, the C/N ratio of ePKA-KTR1.2/tdTomato dropped rapidly, reaching significance within 1 minute (0.8-fold, *p* = 0.005), half-maximum at 3 min (0.7-fold, *p* = 0.0005), and completion at t = 11 minutes (0.5-fold, *p* = 0.005) (***movie S1***). As for cells treated with Fsk and then EGF (10 nM), activation of the EGF/EGFR signaling pathways had no effect on the Fsk-induced C/N ratio of ePKA-KTR1.2/tdTomato.

**Figure 3.**
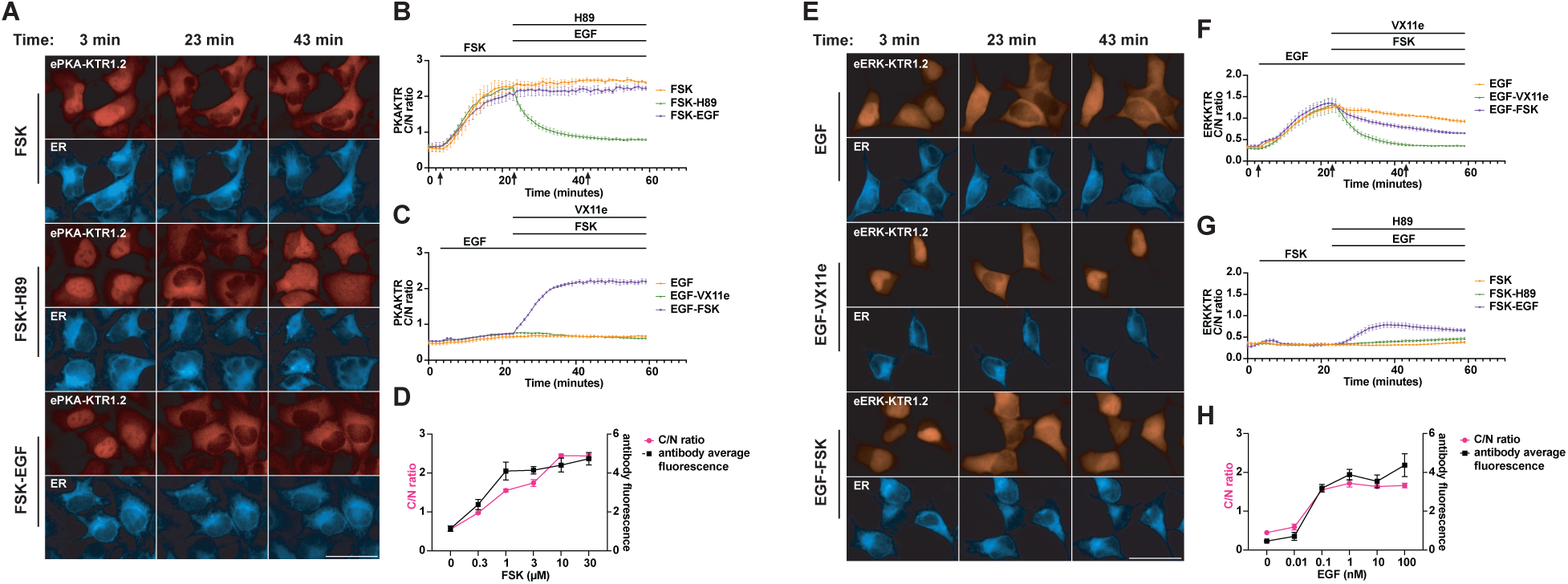
Kinetics of ePKA-KTR1.2 and eERK-KTR1.2 responses. (**A**) Representative fluorescence micrographs of ePKA-KTR1.2/tdCherry-expressing, ER-mTagBFP2-expressing HEK293 cells at t = 3 minutes, t = 23 minutes, and t = 43 minutes, with cells exposed to Fsk alone at t = 3 min, Fsk at t = 3 min and H89 at t = 23 min, or Fsk at t = 3 min and EGF at t = 23 min. Bar, 50 μm. (**B**) Plot of ePKA-KTR1.2/tdCherry C/N ratios at every minute in cells exposed to (orange) Fsk only, (green) Fsk followed by H89, and (purple) Fsk followed by EGF. Lines at top denote times of chemical addition; arrows denote the times at which images in A were taken. (**C**) Plot of ePKA-KTR1.2/tdCherry C/N ratios every minute, in cells exposed to (orange) EGF alone, (green) EGF then VX11e, or (purple) EGF then Fsk. (**D**) Double plot of (red) ePKA-KTR1.2/tdCherry C/N ratios and (black) anti-phosphoPKA antibody staining intensity, following a 30 min incubation in 0, 0.3, 1, 3, 10, and 30 uM Fsk. (**E**) Representative fluorescence micrographs of eERK-KTR1.2/emiRFP670-expressing HEK293 cells at t = 3 minutes, t = 23 minutes, and t = 43 minutes, with cells exposed to EGF alone at t = 3 min, EGF at t = 3 min and VX11e at t = 23 min, or EGF at t = 3 min and Fsk at t = 23 min. Bar, 50 μm. (**F**) Plot of eERK-KTR1.2/emiRFP670 C/N ratios at every minute in cells exposed to (orange) EGF only, (green) EGF then VX11e, and (purple) EGF then Fsk. (**G**) Plot of eERK-KTR1.2/emiRFP670 C/N ratios at every minute in cells exposed to (orange) Fsk only, (green) Fsk then H89, and (purple) Fsk then EGF. (**H**) Double plot of (red) eERK-KTR1.2/emiRFP670 C/N ratios and (black) anti-phospho-ERK antibody staining intensity following a 5 min incubation in 0, 0.01, 0.1, 1, 10, or 100 nM EGF. Data are from a minimum of three independent biological replicates.

In parallel, we also exposed the ePKA-KTR1.2-expressing cell line to EGF alone (10 nM, at t = 3 min). EGF triggered a slight rise in the ePKA-KTR1.2 C/N ratio, and subsequent addition of the ERK inhibitor VX11e (5 µM) at t = 23 appeared to reverse this mild effect, consistent with prior reports that EGF can have a mild activating effect on PKA signaling (*42–45*). Interestingly, when we cells were exposed to EGF at t = 3 and then Fsk (3 µM) at t = 23 min, the responsiveness of ePKA-KTR1.2/tTomato to Fsk reached significance within 1 min (1.2-fold, *p* = 0.0012) and half-maximal at 5 minutes (1.9-fold, p = 0.001) (***Fig. 3C***), raising the possibility that EGF may sensitize the cells to subsequent activators of PKA.

We next tested whether the sensitivity of the ePKA-KTR1.2/tdTomato biosensor was similar to that of the ‘the ‘gold standard’ assay for measuring PKA activity in fixed cells, namely immunostaining with anti-phospho-PKA antibodies (*37, 46–48*). Specifically, HEK293 cells expressing ePKA-KTR1.2/tdTomato were exposed to different concentrations of Fsk for 30 min, after which they were fixed, permeabilized, stained with anti-phospho-PKA antibodies, and interrogated by fluorescence microscopy to measure both p-PKA staining intensity and ePKA-KTR1.2/tdTomato C/N ratio. Digital image analysis was performed on at least ten cells from three independent regions of interest (ROIs), from a minimum of three independent trials. The ePKA-KTR1.2/tdTomato C/N ratios and the anti-phospho-PKA immunostaining intensities were of similar sensitivities and dynamic range (***Fig. 3D***), providing further validation that ePKA-KTR1.2/tdTomato is useful for measuring relative PKA activities.

### Kinetics, specificity, and sensitivity of eERK-KTR1.2

We next interrogated the response kinetics of eERK-KTR1.2. Using an HEK293 cell line stably expressing eERK-KTR1.2/emiRFP670 (∼46 kDa in size) and ER-mTagBFP2, we found that EGF (10 nM, at t = 3 min) triggered an increase in the C/N ratio of eERK-KTR1.2/emiRFP670 that reached significance at 3 minutes (1.5-fold, *p* = 0.002), was half-maximal at 9 min (2.4-fold, *p* = 0.003), and plateaued at 20 min (4-fold, p = 0.0015) (***Fig. 3E***). After reaching this peak, the C/N ratio of eERK-KTR1.2/emiRFP670 declined slowly (0.7-fold between t = 25 and t = 59, *p* = 0.007). In a parallel culture, we followed the addition of EGF at t = 3 min with the ERK inhibitor VX11e (5 µM) at t = 23 min, which triggered a drop in the C/N ratio that reached significance by 8 min (0.5-fold, *p* = 0.049) and appeared complete by 15 min (0.3-fold, p = 0.04) (***Fig. 3E, F***) (***movie S2***).

Previous studies have established that PKA inhibits signaling by the EGF/EGFR/Ras/Raf/ERK pathway by its inhibitory phosphorylation of Raf, and perhaps also by PKA-mediated activation of protein phosphatase 2A(*49–53*). To determine whether eERK-KTR1.2/emiRFP670 can detect this inhibitory cross-talk, we exposed EGF-stimulated cells (10 nM, at t = 3 min) to Fsk at t = 23 min (3 µM), and found that Fsk activation of PKA did indeed trigger the return of eERK-KTR1.2 to the nucleus (***movie S3***). We also found that exposing cells to Fsk at t = 3 min (3 µM) attenuated the response of eERK-KTR1.2/emiRFP670 to subsequent addition of EGF a(10 nM, at = 23 min) (***Fig. 3G***). ERK can also be activated by protein kinase C (**PKC**)(*54–56*), and adding the PKC agonist phorbol myristate acetate (**PMA**)(*57*); 80 nM, at t = 3 min) also triggered a sharp increase in the C/N ratio of eERK-KTR1.2/emiRFP670, and this too was reversed by subsequent addition of VX11e (5 µM) (***fig. S3***).

In addition, when we compared the responsiveness of eERK-KTR1.2 to that of immunostaining with anti-phospho-ERK antibodies, we found that they both had similar sensitivities to EGF. This is shown here for cells expressing eERK-KTR1.2/emiRFP670 that were exposed to different concentrations of EGF for 5 min, then fixed, permeabilized, stained with anti-phospho-ERK antibodies, and interrogated by fluorescence microscopy (***Fig. 3D***).

### Simultaneous live cell imaging of PKA activity, ERK activity, and calcium

To determine whether co-expression of these eKTRs allowed the simultaneous monitoring of PKA and ERK activities in real-time, we created an HEK293 cell line that stably co-expressed ePKA-KTR1.2/tdTomato, eERK-KTR1.2/emiRFP670, and ER/mTagBFP2. Addition of Fsk triggered the nuclear export of ePKA-KTR1.2/tdTomato and a slight drop in the eERK-KTR1.2 C/N ratio, subsequent addition of EGF led to an intermediate rise in the eERK-KTR1.2/emiRFP670 C/N ratio, addition of H89 triggered the nuclear import of ePKA-KTR1.2, and later addition of VX11e had no effect on either sensor (Fig. 4A, B) (***movies S4A (ePKA-KTR1.2) and S4B (eERK-KTR1.2***)). In another set of experiments, we found that EGF induced a sharp rise in the C/N ratio of eERK-KTR1.2, and once again also induced a shallow rise the ePKA-KTR1.2 C/N ratio, consistent with previous reports of EGF-triggered activation of PKA(*42–45*). Subsequent addition of Fsk drove the ePKA-KTR1.2 C/N ratio even higher, while later addition of VX11e reduced the eERK-KTR1.2 C/N ratio, with subsequent addition of H89 dropping the C/N ratio of ePKA-KTR1.2 (***Fig. 4C, D***) (***movies S5A (ePKA-KTR1.2) and S5B (eERK-KTR1.2***)).

**Figure 4.**
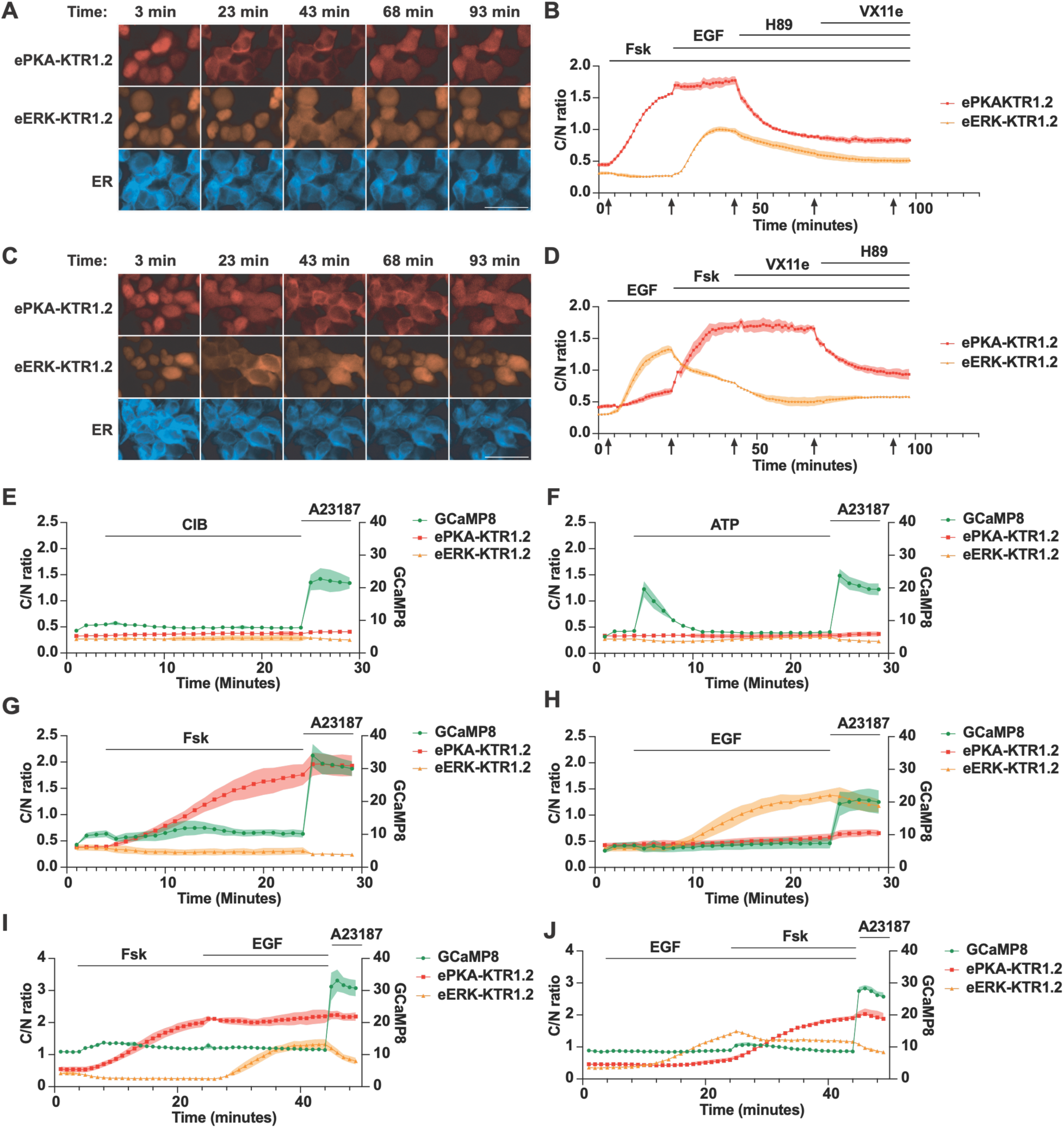
Simultaneous measurement of PKA activity, ERK activity, and calcium. (**A**) Representative fluorescence images of HEK293 cells co-expressing ePKA-KTR1.2/tdTomato, eERK-KTR1.2/emiRFP670, and ER-mTagBFP2 exposed to Fsk at t = 3 min, then EGF at t = 23 min, then H89 at t = 50 min, then VX11e at t = 75 min, with images collected at t = 3, 23, 43, 68, and 93 min. Bar, 50 μm. (**B**) Plot of C/N ratios of (red) ePKA-KTR1.2/tdTomato and (orange) eERK-KTR1.2/emiRFP670 every 60 seconds, showing their responses to Fsk, then EGF, then H89, then VX11e. Lines at the top denote the times of chemical addition, and arrows at the bottom denote the times at which the images of A were collected. (**C**) Representative fluorescence images of HEK293 cells co-expressing ePKA-KTR1.2/tdTomato, eERK-KTR1.2/emiRFP670, and ER-mTagBFP2 exposed to Fsk at t = 3 min, then EGF at t = 23 min, then H89 at t = 50 min, then VX11e at t = 75 min. Bar, 50 μm. (**D**) Plot of C/N ratios of (red) ePKA-KTR1.2/tdTomato and (orange) eERK-KTR1.2/emiRFP670 every 60 seconds, showing their responses to EGF, then Fsk, then VX11e, then H89. (**E-J**) Plots of (green) GCaMP8 fluorescence intensity, (red) ePKA-KTR1.2/tdTomato C/N ratio, and (dark red) eERK-KTR1.2/emiRFP670 C/N ratio calculated from images collected at every minute as the cells were exposed to (E) CIB then A23187, (F) ATP then A23187, (G) Fsk then A23187, (H) EGF then A23187, (I) Fsk then EGF then A23187, or (J) EGF then Fsk then A23187. Lines at the top denote the times of chemical addition, C/N ratios are reflected in the left Y-axis, and GCaMP fluorescence intensity is reflected in the right Y-axis. Data are from a minimum of three independent biological replicates.

We also multiplexed these PKA and ERK eKTRs with the genetically-encoded calcium indicator GCaMP8s(*58*). A GCaMP8s expression vector was transfected into the cell line that expresses ePKA-KTR1.2/tdTomato, eERK-KTR1.2/emiRFP670, and ER-mTagBFP2, and the co-expressing cells were examined by fluorescence microscopy as they were exposed to modulators of calcium, PKA, and ERK. During cell incubation in calcium imaging buffer (CIB), further addition of CIB had no impact on any of these sensors. The calcium ionophore. A23187 (200 μM) triggered an immediate leap in GCaMP fluorescence intensity (***Fig. 4E***), whereas exposure to ATP (10 μM) triggered a transient rise in GCaMP fluorescence, presumably via P2Y receptor-triggered, store-operated intracellular calcium release(*59*) (***Fig. 4F***). In contrast, elevated calcium did not affect either the PKA or ERK eKTR C/N ratios. Each eKTR did, however, respond as expected to its cognate activator (***Fig. 4G-J***). In what might be a sign of PKA-calcium cross signaling, we found that Fsk led to a mild increase in GCaMP8s fluorescence intensity.

### The ePKA-KTR1.4 and eERK-KTR1.2 variants are appropriately tuned for DRG neurons

Given the wide array of specialized functions performed by distinct cell types, its possible that the detection of PKA and ERK activities in a single cell type may require eKTRs tuned to its particular kinase activities and nuclear import-export dynamics. To explore this possibility, we examined eKTR performance in primary sensory neurons, which rely heavily on PKA and ERK signaling pathways(*60, 61*). We dissociated and plated mouse dorsal root ganglion neurons, transduced them with ePKA-KTR1.2/tdTomato-encoding lentivirus, incubated them for 3 days to allow for KTR expression, and then monitored ePKA-KTR1.2/tdTomato subcellular distribution in response to Fsk and H89. In brief, we added Fsk at t = 0 (3 µM), added the PKA inhibitor H89 at t = 15 min (50 µM), while capturing images at t = 0, t = 15 min, and t = 45 min. These experiments demonstrated that the C/N ratio of ePKA-KTR1.2/tdTomato rose significantly in response to Fsk and returned to baseline in response to H89 (***Fig. 5A***). However, these experiments also showed that the C/N ratio of ePKA-KTR1.2/tdTomato was substantially higher than 1 in DRG neurons prior to any stimulation, and as a result, its dynamic range in these cells was only 1.5-fold (C/N ratio rising from ∼1.7 to 2.2), which is far from optimal for a KTR. Fortunately, our earlier work had also developed the ePKA-KTR1.4 sensor, which has a stronger NLS. When expressed in DRG neurons, ePKA-KTR1.4/tdTomato had a much lower resting C/N ratio, ∼0.8, compared to ePKA-KTR1.2/tdTomato (***Fig. 5A***). Even more importantly, the ePKA-KTR1.4/tdTomato C/N ratio rose to ∼2.4 in response to Fsk, demonstrating a 3-fold dynamic range, and returned to baseline in response to H89, demonstrating its sensitivity to PKA agonists and inhibitors.

**Figure 5.**
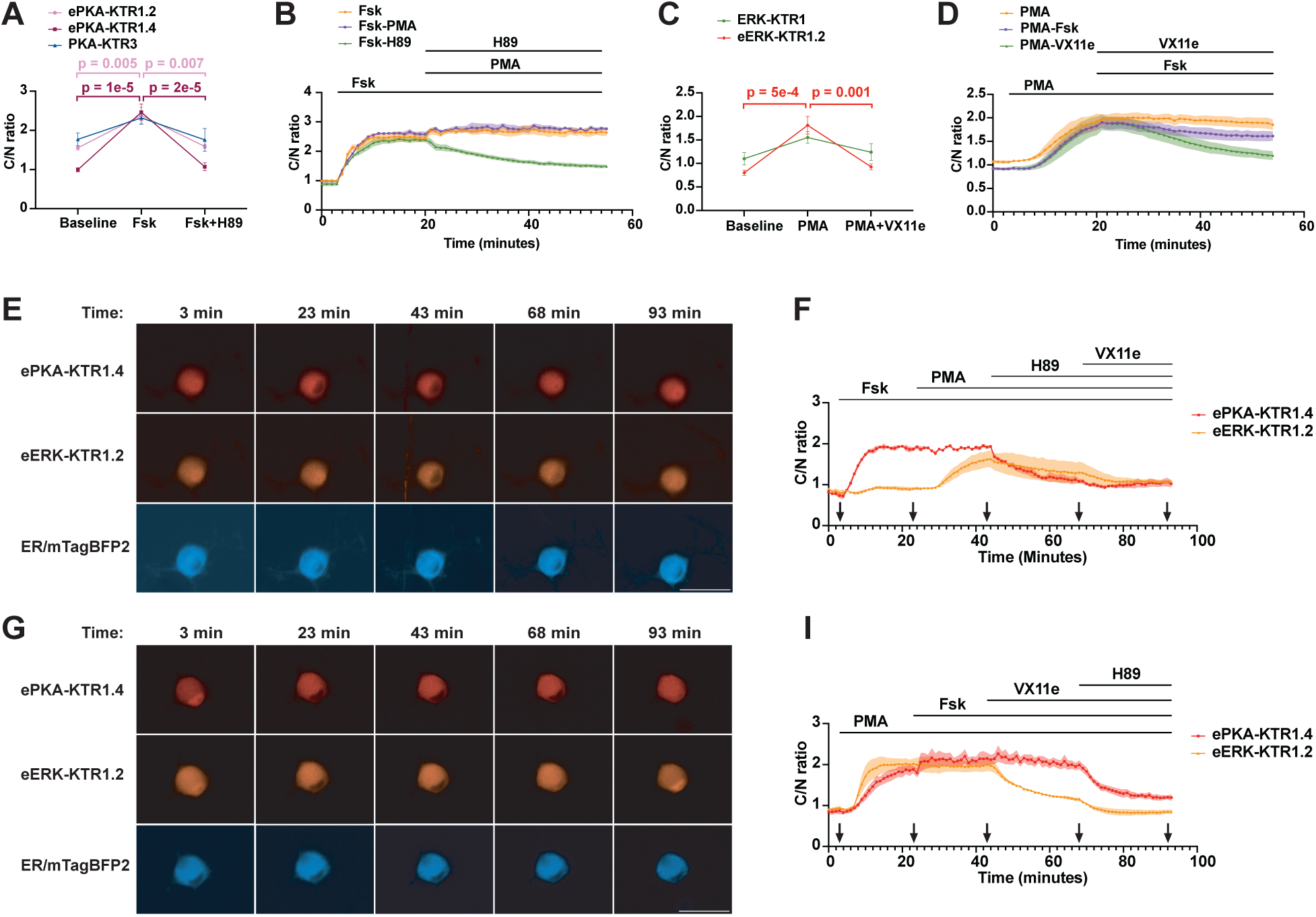
Superior performance of ePKA-KTR1.4 and eERK-KTR1.2 in DRG neurons. (**A**) Graph of C/N ratios of (pink) ePKA-KTR1.2, (purple) ePKA-KTR1.4, and (green) PKA-KTR3 in DRG neurons, at baseline, in response to Fsk, and in response to subsequent addition of H89. ** and **** denote *p* values <0.01 or <0.0001, respectively. The *p* values for the changes in PKA-KTR3 C/N ratio were 0.07 and 0.06, respectively. (**B**) Plot of C/N ratios of ePKA-KTR1.4/tdTomato collected every minute in response to (orange) Fsk alone, (green) Fsk followed later by addition of H89, or (purple) Fsk followed later by addition of PMA. Lines at the top denote the times of chemical addition. (**C**) Graph of C/N ratios of (green) ERK-KTR1 and (orange) eERK-KTR1.2 in DRG neurons at baseline, in response to PMA, and in response to subsequent addition of H89. *** denote *p* values <0.001. The *p* values for the changes in ERK-KTR1 C/N ratio were 0.08 and 0.3, respectively. (**D**) Plot of C/N ratios of eERK-KTR1.2/emiRFP670 collected every minute in response to (orange) PMA alone, (green) PMA then VX11e, or (purple) PMA then VX11e. Lines at the top denote the times of chemical addition. (**E**) Representative fluorescence images of DRG co-expressing ePKA-KTR1.4/tdTomato, eERK-KTR1.2/emiRFP670, and ER-mTagBFP2, exposed sequentially to Fsk at t = 3 min, then PMA at t = 23 min, then H89 at t = 45 min, then VX11e at t = 70 min. Images were collected at t = 3, 23, 43, 68, and 93 minutes. Bar, 50 μm. (**F**) Plot of C/N ratios of (red) ePKA-KTR1.4/tdTomato and (orange) eERK-KTR1.2/emiRFP670 at 1-minute intervals, showing their responses to Fsk, then PMA, then H89, then VX11e. (**G**) Representative fluorescence images of DRG co-expressing ePKA-KTR1.4/tdTomato, eERK-KTR1.2/emiRFP670, and ER-mTagBFP2, exposed sequentially to PMA at t = 3 min, then Fsk at t = 23 min, then VX11e at t = 45 min, then H89 at t = 70 min. Images were collected at t = 3, 23, 43, 68, and 93 minutes. Bar, 50 μm. (**H**) Plot of C/N ratios of (red) ePKA-KTR1.4/tdTomato and (orange) eERK-KTR1.2/emiRFP670 at 1-minute intervals, showing their responses to PMA, then Fsk, then VX11e, then H89. Data are from a minimum of three independent biological replicates. *p* values were calculated by two-way ANOVA.

During the course of our studies, two additional PKA KTRs were reported, which we refer to as PKA-KTR2(*28*) and PKA-KTR3(*29*). To determine whether PKA-KTR2 or PKA-KTR3 displayed sensitivities similar to, or better, than the ePKA-KTRs described earlier in this study, we fused their sensor domains to the N-terminus of mCherry and tdCherry, expressed them in HEK293 cells, and monitored their C/N ratio in response to Fsk. Neither PKA-KTR2/mCherry nor PKA-KTR2/tdCherry displayed a significant change in response to Fsk (***fig. S4A, B***), indicating that its sensor domain does not function in the context of these proteins. As for PKA-KTR3, we found that the C/N ratio of PKA-KTR3/mCherry rose significantly in response to Fsk, but did not exceed 1 (***fig. S4C***), indicating that this particular protein has only a limited ability to sense PKA activity. In contrast, we found that PKA-KTR3/tdCherry displayed a much larger Fsk-induced increase in its C/N ratio, ∼3-fold (***fig. S4D***), consistent with our hypothesis that KTRs work better when they’re too large to rapidly diffuse through the nuclear pore. In light of these results, we tested whether PKA-KTR3/tdCherry was able to report PKA activity in DRG neurons. However, when we expressed PKA-KTR3/tdCherry in DRG neurons we found that its resting C/N ratio was extremely high, ∼1.8, and that it failed to respond significantly to Fsk or to subsequent addition of H89 (***Fig. 5A***).

Given its superior performance characteristics in DRG neurons, we next examined the kinetics of ePKA-KTR1.4/tdTomato responses in these primary sensory neurons. Transduced neurons exposed to Fsk (3 µM) at t = 3 min. led to a sharp increase in its C/N ratio, which remained high over the course of the experiment (***Fig. 5B***). In a parallel population of cells stimulated with Fsk (3 µM), and then later exposed to the PKA inhibitor H89 (50 µM at t = 20 min.), we observed that H89 led to a drop in the ePKA-KTR1.4/tdTomato C/N ratio. In Fsk-induced neurons, subsequent addition of PMA had no effect.

We next tested which KTR was best suited to monitor ERK activity in DRG neurons. Specifically, we transduced DRG neurons with lentiviruses designed to express either ERK-KTR1/emiRFP670, which has a weaker NLS, or eERK-KTR1.2/emiRFP670, which has a stronger NLS. We then examined the neurons by fluorescence microscopy at baseline, in response to ERK activation by the PKC agonist PMA (80 nM) at t = 0 min. and in response to the ERK inhibitor VX11e at t = 15 min (5 µM), with images collected at t = 0, t =15 min, and t = 45 min. (***Fig. 5C***). eERK-KTR1.2/emiRFP670 responded significantly to PMA, shown here by the rise in its C/N ratio from ∼1 to ∼2, and also responded significantly to VX11e, which caused its C/N ratio to return to baseline. In contrast, ERK-KTR1/emiRFP670 did not respond significantly to either PMA or VX11e.

The ability of ePKA-KTR1.4/tdTomato and eERK-KTR1.2/emiRFP670 to report PKA and ERK activities in DRG neurons was also examined in co-transduced neurons. In one set of experiments, co-expressing neurons were exposed serially to Fsk, PMA, H89, and VX11e. We found that Fsk-induced the cytoplasmic translocation of ePKA-KTR1.4/tdTomato, and also attenuated the PMA-induced increase in eERK-KTR1.2/emiRFP670 C/N ratio, while H89-mediated the nuclear import of ePKA-KTR1.4/tdTomato, and VX11e-mediated the re-import of eERK-KTR1.2/emiRFP670 (***Fig. 5E, F***). In a second set of experiments, co-expressing DRG neurons exposed serially to PMA, Fsk, VX11e, and H89 revealed the expected PMA-mediated nuclear export of eERK-KTR1.2/emiRFP670, but also exhibited a strong PMA-mediated nuclear export of ePKA-KTR1.4/tdTomato, Subsequent responses to Fsk, VX11e, and H89 were otherwise as expected (***Fig. 5 G, H***). The PMA-mediated increase in PKA signaling seen in DRG neurons was also apparent in PMA-treated HEK293 cells (***fig. S5***), and might reflect the previously reported activation of PKA by EGF/ERK signaling(*42–45*).

## DISCUSSION

Among the integrative kinase biosensors, KTRs have the fastest response kinetics, reporting changes on a minute-by-minute timescale, similar to the kinetics of many kinase-regulated physiological processes. As such, KTRs fill an important temporal niche, recording kinase activities over periods of time that are ∼100-times faster than transcriptional reporters and conditionally stabilized reporters, which record kinase activities over hours-to-days(*14–23*), though they are ∼10-100 times slower than the fast-responding FRET and ratiometic biosensors. As a result, KTRs provide a unique readout of kinase activities integrated over the timescale of many biological processes. Although the ability to monitor integrated kinase activities on a minute-by-minute basis is key to understanding PKA, ERK, and other kinase signaling pathways in many biological contexts, there has been little if any development of enhanced KTRs with greater sensitivity and dynamic range has been limited(*35*), especially in comparison to the tremendous improvements in fast-response kinase biosensors(*13, 58*).

The data presented here show that KTRs can be improved considerably by careful consideration and optimization of key variables that affect their (1) passive diffusion through the nuclear pore and (2) active nuclear import and export. For example, the operating hypothesis underlying all KTRs is that their their passive diffusion through the nuclear pore should be kept to a minimum, so that their relative distribution between nucleus and cytoplasm reflects the dynamic balance between their NLS-mediated nuclear import and NES-mediated nuclear export. Not surprisingly, increasing the size of KTRs so they could no longer diffuse quickly through the nuclear pore complex allowed us to identify the primary reason why the original PKA-KTR described by Regot et al.(*24*) didn’t work particularly well, namely that its’ NLS was simply too weak to drive nuclear protein import. This, in turn, allowed us to design enhanced PKA sensor domains that contained NLSs of different strength. Together, varying KTR size and NLS strength allowed us to convert the relatively unresponsive PKA-KTR1(*24*) into the highly responsive ePKA-KTR1.2 and ePKA-KTR1.4 biosensors, to convert ERK-KTR1(*24*) into the more nuclear eERK-KTR1.2 sensor, and to show that these enhanced KTRs are required for monitoring PKA and ERK activities in primary sensory neurons.

Our studies also showed the power of KTRs for simultaneous monitoring of PKA and ERK activities, as multiplexed expression of ePKA-KTR1.2/tdTomato, eERK-KTR1.2/emiRFP670, ER/mTagBFP, and the calcium sensor GCaMP8s allowed us to simultaneously monitor the PKA activity, ERK activity, and calcium levels in individual live cells, in response to known agonists and inhibitors of PKA, ERK, calcium, and even PKC. These crosstalk effects have been described previously, including PKA-mediated inhibition of ERK(*49–53*), PMA/PKC-mediated activation of ERK(*56*), the mild EGF-mediated activation of PKA(*42–45*), and the strong activation of PKA by PMA, which we observed in neurons but not HEK293 cells. Taken together, these results lend further support to the idea that the enhanced ePKA-KTRs and eERK-KTRs are useful tools for monitoring these complex and interactive kinase signaling pathways. Our results also point to a potential effect of PKA activation on intracellular calcium levels, as Fsk in all cases induced a slight rise in GCMP8s fluorescence intensity. It will be interesting to see whether further improvements in KTR performance characteristics might be achieved by altering other variables likely to impact KTR dynamics (e.g. increasing sensor domain copy number, KTR oligomerization, sensor domain position, FP identity, etc.), and whether the new rules for KTR design established in this study can be used to improve the dynamic range and sensitivities of other KTRs for other kinases.

## METHODS

### Cell culture and DNA transfections

HEK293 cells and HEK293T cells (ATCC CRL-1573 and ATCC CRL-3216, respectively) were grown in complete medium (**CM**: Dulbecco’s Modified Eagle’s Media (ThermoFisher, 11965092) supplemented with 10% fetal bovine serum (Gibco, 16000044) and 1% penicillin/streptomycin (Gibco, 15140122)). Cultures were grown in 90% humidity and 5% CO2. Cells were transfected by lipofection using Lipofectamine 3000 according to the manufacturer’s instructions (Thermo, L3000015). Excised DRG neurons were cultured in neuron culture media, consisting of Neurobasal Plus (Gibco, A3582901) supplemented with B27 (Gibco, 17504044).

### Plasmids

Plasmids were assembled by ligation of restriction enzyme-cleaved DNA fragments, which were derived from other plasmids or for PCR-amplified synthetic DNAs (geneblocks, IDT Inc.). Plasmid architectures were confirmed by combination of restriction enzyme mapping and DNA sequencing, with all sequences derived from amplified fragments confirmed in their entirety by DNA sequence analysis. All KTR vectors, mNeonGreen vectors, and mTagBFP2 vectors are based on Gould lab pC or pLenti vectors(*62*). Plasmid descriptions, including deduced amino acid sequences of key ORF, are presented in the supplemental information (***table S1***). Addgene was the source of the plasmids psPAX2 from Didier Trono (Addgene# 12260) and VSV.G from Tannishtha Reya (Addgene# 14888).

### Fluorescence microscopy

Live-cell fluorescence widefield microscopy was carried out using the EVOS M7000 Imaging System (Thermo Fisher, AMF7000) and 20x objective, equipped with an EVOS Onstage Incubator (Thermo Fisher, AMC2000) set to 37 degrees Celsius. Prior to imaging, growth medium was replaced with HEPES-buffered physiological imaging media (10 mM HEPES, 130 mM NaCl, 3 mM KCl, 2.5 mM CaCl_2_, 0.6 mM MgCl_2_, NaHCO_3_, 10 mM Glucose). Images of HEK293 cells were captured by automated robotic imaging of four independent fields of view, while images of DRG neurons were captured from 10 independent fields of view.

### Image analysis and quantification of C/N ratios

Images were interrogated using NIS Elements software (RRID:SCR_014329) version 4.60. For endpoint multicolor image analyses, a pair of regions of interests (ROIs) representing a randomly selected area in the nucleus and cytoplasm for each cell were drawn based on either a combination of H2B-mTagBFP2 and mNeonGreen fluorescence distributions in experiments of Fig. 1 and 2, or ER-mTagBFP2 distribution alone in all subsequent experiments. For each image, a minimum of 10 cells were analyzed. KTR fluorescence intensity in the cytoplasmic and nuclear compartments was calculated from the average pixel fluorescence intensity within each compartment ROI, and the cytoplasm/nucleus fluorescence ratio (C/N ratio) was calculated for each cell, followed by averaging across all assayed cells in each image to create region-of-interest data points, which were then treated as individual data points across a minimum of three biological replicates. Calculations of mean, standard error of the mean (s.e.m.), and ANOVA p values were carried out using GraphPad Prism. Image analyses were conducted in a double-blinded fashion. For time course analyses, multicolor images were first registered in Fiji (ImageJ), and exported as movies. Analytical procedures were similar to those described above, except that the positions of cytosolic and nuclear ROIs in each cell were checked between the first and last frame of the movie (using reference marker channels), to account for mispositioning over time due to cell movement over the course of the experiment. The Python script for KTR analysis is available

### Agonists, inhibitors, and antibodies

A23187 (Sigma, C7522), ATP (Sigma, A26209), Forskolin (Cayman Chemical, 11018), H89 (MedChemExpress, HY-15979A), EGF (Gibco, PHG0311L), VX-11e (Selleckchem, S7709), leptomycin B (Sigma Aldrich, L2913), Phorbol 12-myristate 13-acetate (Sigma Aldrich, P1585) were obtained from commercial sources. Anti-phospho-ERK antibody (used at 1:1000; Cell Signaling Technology, 9101), anti-phospho-PKA antibody (used at 1:1000; Abcam, ab32390), and fluorophore-conjugated secondary antibodies (used at 1:800; Jackson ImmunoResearch) were also obtained from commercial sources.

### Lentivirus production

HEK293T cells were seeded into 6-well plates the day prior to transfection in a volume of 2 mL complete medium per well. The next day (d0), the cells were at ∼90% confluency and transfected with equal amounts (by mole) of the pLenti transfer vector (designed to express the KTR-2a-ER-mTagBFP2 ORF), psPAX2, and VSV.G. The next day (d1), the 2mL culture medium was collected at place in a 50 mL sterile centrifuge tube, which was used as the collection vessel for all conditioned medium samples over the course of the lentivirus preparation. Cells from each well were released by trypsinization and re-seeded on a 10 cm plate with 15 mL complete medium (CM). The next day (d2), 10 mL of supernatant was collected and added to the 2 mL of conditioned medium collected the previous day, and 10 mL of fresh cell culture media was added to each 10 cm dish of cells and the cells were incubated overnight. The next day (d3), 10 mL of supernatant was collected from each dish and added to the 12 mL of conditioned medium collected over the previous two days, and 10 mL of fresh cell culture media was added to each 10 cm dish of cells and the cells were again incubated overnight. The next day (d4), all 15 mL of conditioned medium was collected and added to the 22 mL of conditioned medium that had been collected from each sample over the previous 3 days, yielding a total volume of 37 mL conditioned medium from transfected cell population. The 37 mL was passed through a 450 nm pore size, syringe tip filter unit, yielding 37 mL of clarified conditioned medium. Lentivirus particles were pelleted from these samples by ultracentrifugation at 100,000x *g* for 120 minutes (Optima L-90K, using a SW-32 Ti rotor, Beckman), and the pellets were resuspended in either 1mL of complete medium or neuron culture media.

### Lentiviral transductions

For transduction of HEK293 cells, were seeded into 24-well tissue culture plates at 10% confluency, grown overnight in complete medium, and then incubated with 50-100 uL of lentivirus preparation. Cells were then incubated for 2-4 more days prior to use in imaging or fluorescence activated cell sorting (**FACS**). For transduction of DRG neurons, 50 microliters of lentivirus prepartion was added to primary DRG neurons cultured in 300 microliters of neuron culture medium. Live-cell imaging experiments were conducted 2 to 3 days after transduction.

### Fluorescence activated cell sorting (FACS)

Lentivirus-transduced, KTR-expressing HEK293 cells were cultured for a minimum of 7 days post-transduction, after which they were trypsinized, together with the parental HEK293 cell line, and interrogated by flow cytometry using a Sony MA900 Cell Sorter. Flow cytometry confirmed that HEK293 cells displayed only background fluorescence, that some of the cells transduced with the ePKA-KTR1.2/tdTomato-2a-ER-mTagBFP2-expressing lentivirus displayed bright tdTomato and mTagBFP2 fluorescence, that some of the cells transduced with the eERK-KTR1.2/emiRFP670-2a-ER-mTagBFP2-expressing lentivirus displayed bright emiRFP670 and mTagBFP2 fluorescence, and that some of the cells co-transduced with both lentiviruses expressed bright tdTomato, emiRFP670, and mTagBFP2 fluorescence. We then used the Sony MA900 Cell Sorter to isolate subpopulations of these three fluorescent cell lines that displayed the highest 25% of fluorescence brightness, after which the three FACS-sorted cell lines were recovered in complete medium, expanded, and stocked.

### Mice, DRG isolation, culture, and transduction

All animal studies were approved by the Johns Hopkins University Animal Care and Use Committee (IACUC approval #MO23M18). C57BL/6 mice were maintained under standard conditions, with water and food provided ad libitum. To obtain DRG neurons, C57BL/6 mice of 4-6 weeks old were euthanized by CO_2_ inhalation followed by cervical dislocation prior to dissection. Under a dissection microscope, the spinal column was excised, bisected, and peeled to reveal dorsal root ganglia. Individual ganglia were picked using surgical forceps and placed in ice-cold PBS. Enzymatic digestion of connective tissue was carried out at 37°C using Liberase TM (Roche, 5401119001) for 20 min, followed by Liberase TL (Roche, 5401020001) and papain enzyme suspension (Worthington, LS003126) for 20 min. Digestion was terminated by 2% bovine serum albumin (Sigma, A9418) in neuron culture media. Individual DRG neurons were released by repeated pipetting (20 times) of digested ganglia in 500 µL neuron culture media, which was performed up to 5 times, or until tissue dissociation was complete. Cells were pelleted at 300 x *g* for 3 minutes, resuspended in neuron culture media, and plated onto 8-mm cover glasses (Electron Microscopy Sciences, 7229608) or imaging chambers (Ibidi, 80807) pre-coated with 0.01% poly-D-lysine (Sigma Aldrich, P6407) and mouse laminin (Gibco, 23017015).

### Formal analysis

Experiments were performed a minimum of three times, unblinded, and the statistical interrogation of differences was performe by two-way ANOVA.

## Acknowledgments

The authors thank James Morrell for his outstanding technical assistance, to Dr. Xinhong Dong for use of their FACS machine, and members of the Gould lab, Caterina lab and other colleagues at Johns Hopkins University, especially Dr. Michael Wolfgang, Dr. Sergi Regot and Dr. Takanari Inoue, for their helpful suggestions over the course of these studies. In addition, we thank Mr. Wei Zhou for his development of a comprehensive Python script that we used for KTR analysis.

## Author contributions

Conceptualization: SJT, YG, MJC, SJG Methodology: SJT, YG, FZ, JX, AG, MJC, SJG Investigation: SJT, YG, AD, FZ, JX, AG, MJC, SJG Visualization: SJT, YG, AD, FZ, JX, MJC, SJG Funding acquisition: MJC, SJG Project administration: MJC, SJG Supervision: SJT, MJC, SJG Writing – original draft: SJG Writing – review & editing: SJT, YG, AD, FZ, JX, AG, MJC, SJG

## Data and materials availability

All data are available in the main text or the supplementary materials. All materials are available upon request, provided that a material transfer agreement is completed with Johns Hopkins University.

## SUPPLEMENTAL INFORMATION

**Figure S1.**
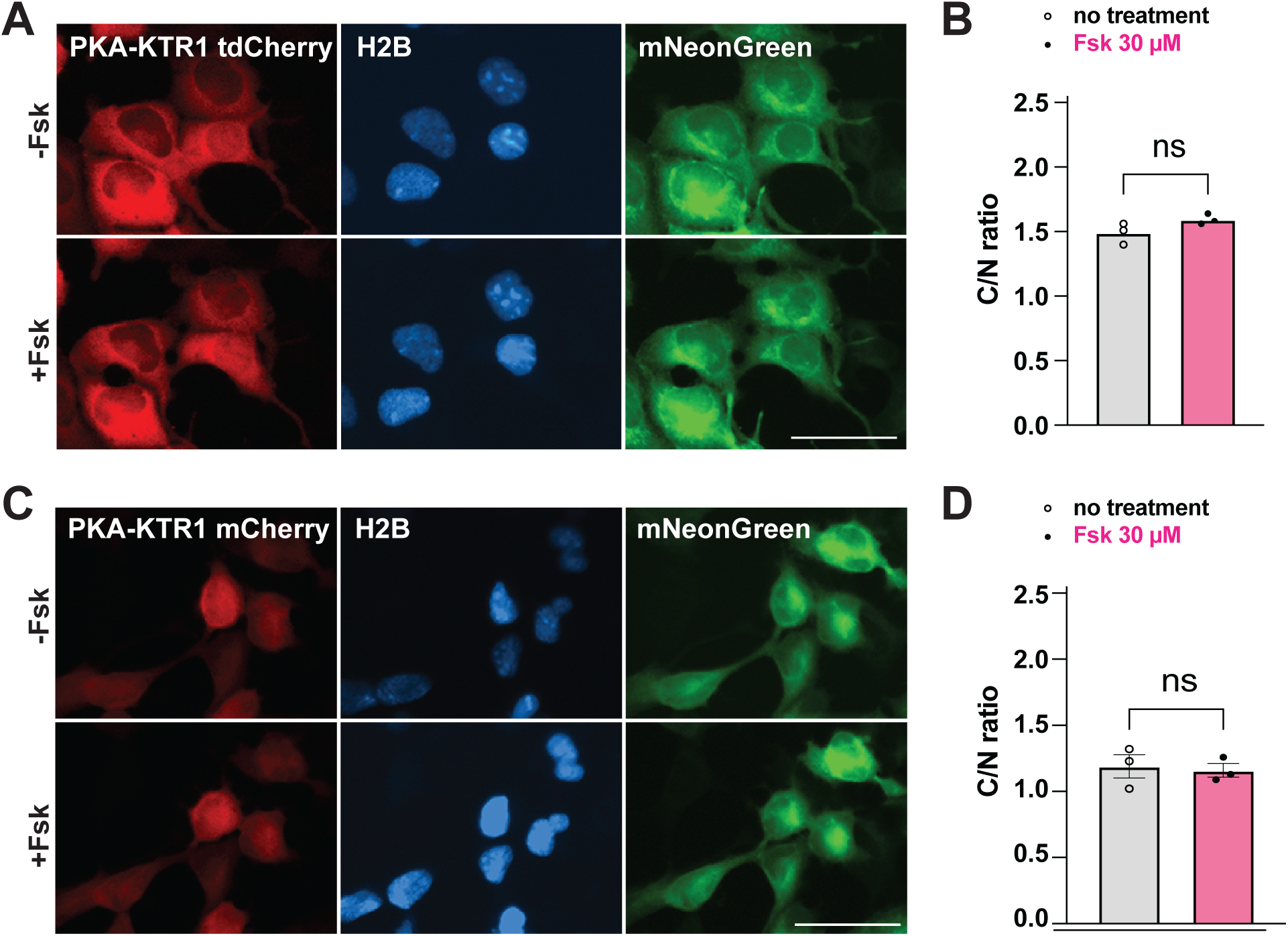
The PKA-KTR1 sensor domain does not drive Fsk-induced protein translocation to the cytoplasm. HEK293 cells were transfected with plasmids designed to co-express H2B-mTagBFP2, mNeonGreen, and (A, B) PKA-KTR1/tdCherry (∼64 kDa), or (C, D) PKA-KTR1-mCherry (∼32 kDa). (**A**) Fluorescence micrographs of HEK293 expressing cells expressing PKA-KTR1/tdCherry before and after 30 minutes incubation in 30 μM Fsk. Bar, 50 μM. (**B**) Bar plot of PKA-KTR1/tdCherry C/N ratio before and after addition of after 30 minutes incubation in 30 μM Fsk. (**C**) Fluorescence micrographs of HEK293 expressing cells expressing PKA-KTR1/mCherry before and after 30 minutes incubation in 30 μM Fsk. Bar, 50 μM. (**D**) Bar plot of PKA-KTR1/mCherry C/N ratio before and after addition of after 30 minutes incubation in 30 μM Fsk. Data are from a minimum of three independent biological replicates.

**Figure S2.**
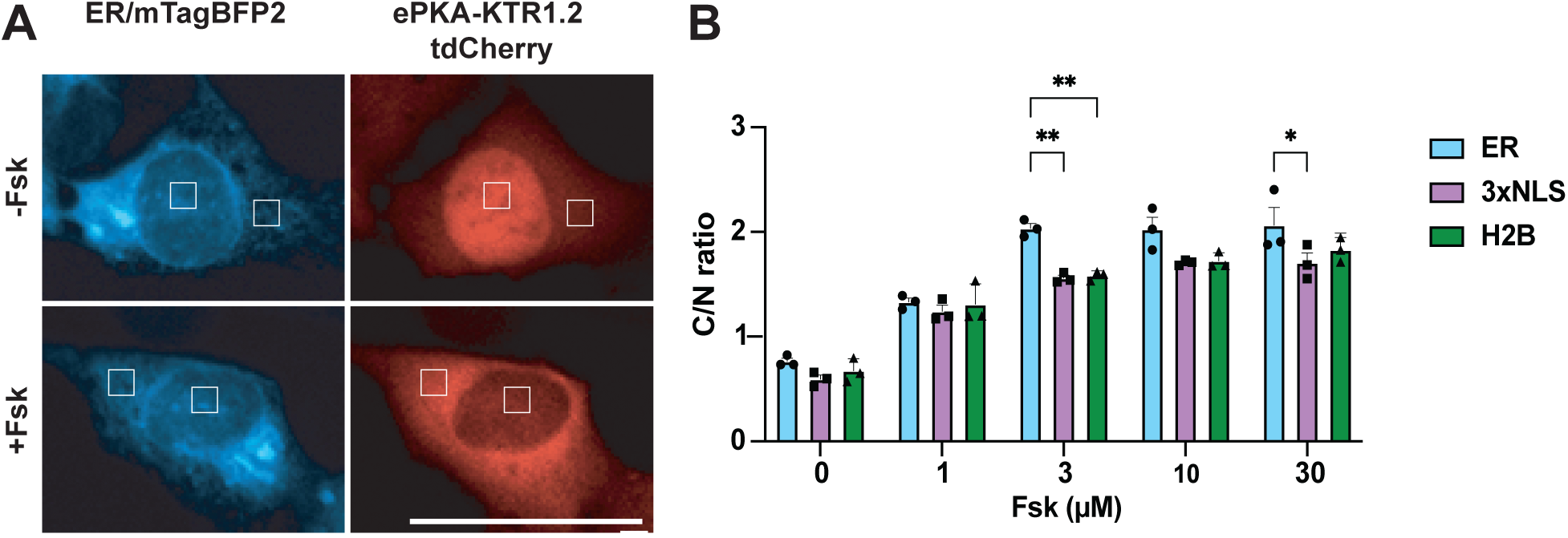
Marking nucleus and cytoplasm with ER-mTagBFP2. (**A**) Fluorescence micrographs of HEK293 cells co-expressing ER-mTagBFP2 and ePKA-KTR1.2/tdCherry. The distribution of ER-mTagBFP2 determines the positioning of ‘counting areas’ (white boxes) that are then used to count the relative fluorescence intensity of KTR proteins in the nucleus and cytoplasm, with the C/N ratio determined from the average fluorescence brightness of all pixels within each counting area. (**B)** C/N ratios of ePKA-KTR1.2 in HEK293 cells co-expressing (blue bars) the ER-mTagBFP2 marker (ER), (purple bars) the 3xNLS-mTagBFP2 marker (3xNLS), or (green bars) the H2B-mTagBFP2 marker (H2B). Cells were exposed to 0, 1, 3, 10, or 30 uM Fsk for 30 minutes, imaged by fluorescence microscopy, and C/N ratios were calculated from digital images. Data are from a minimum of three independent biological replicates. ANOVA *p* values are denoted by * <0.05, and ** <0.01.

**Figure S3.**
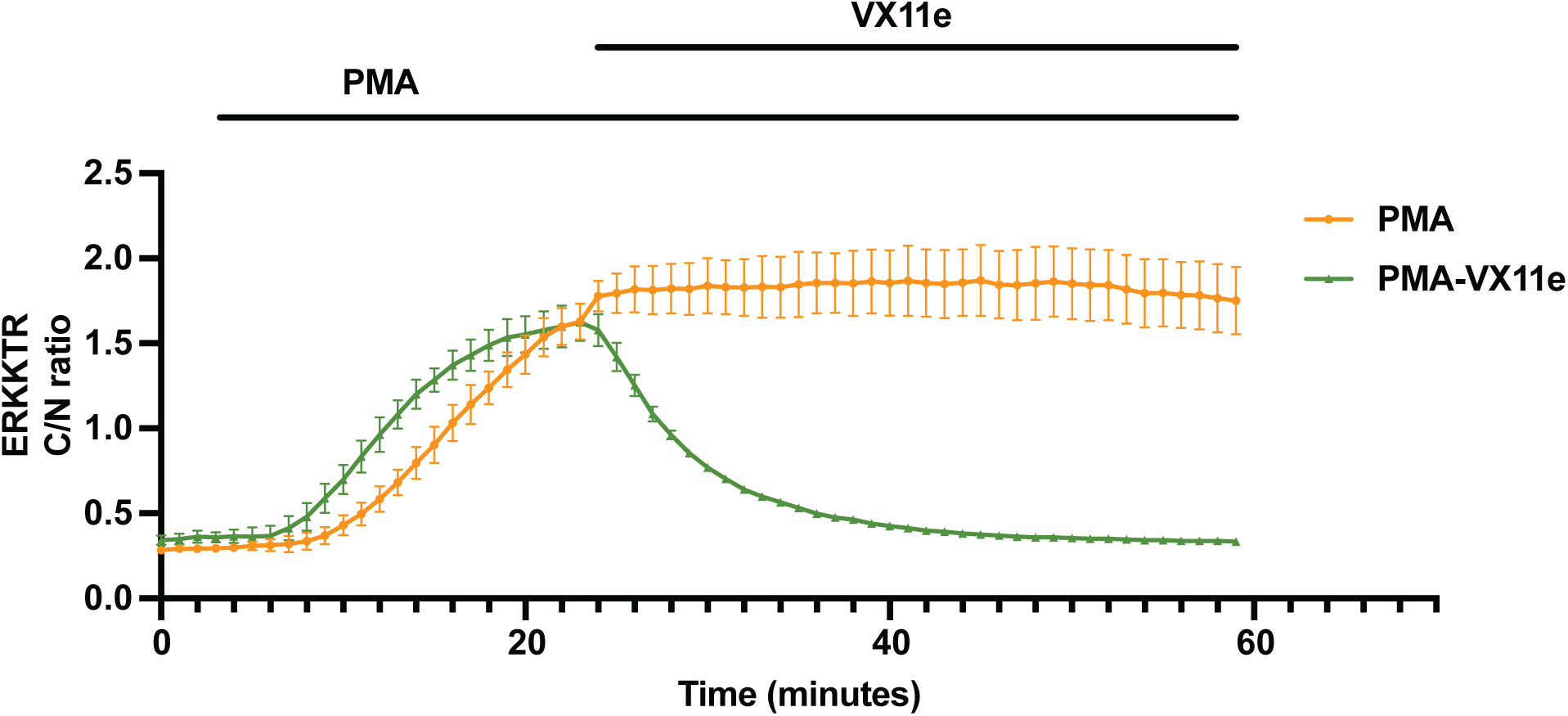
Response of eERK-KTR1.2/emiRFP670 to PMA, and to PMA then VX11e. HEK293 cells expressing eERK-KTR1.2/emiRFP670 were imaged every 60 sec (orange) in response to PMA alone or (green) in response to PMA, followed 20 min later by addition of VX11e. Data are from a minimum of three independent biological replicates.

**Figure S4.**
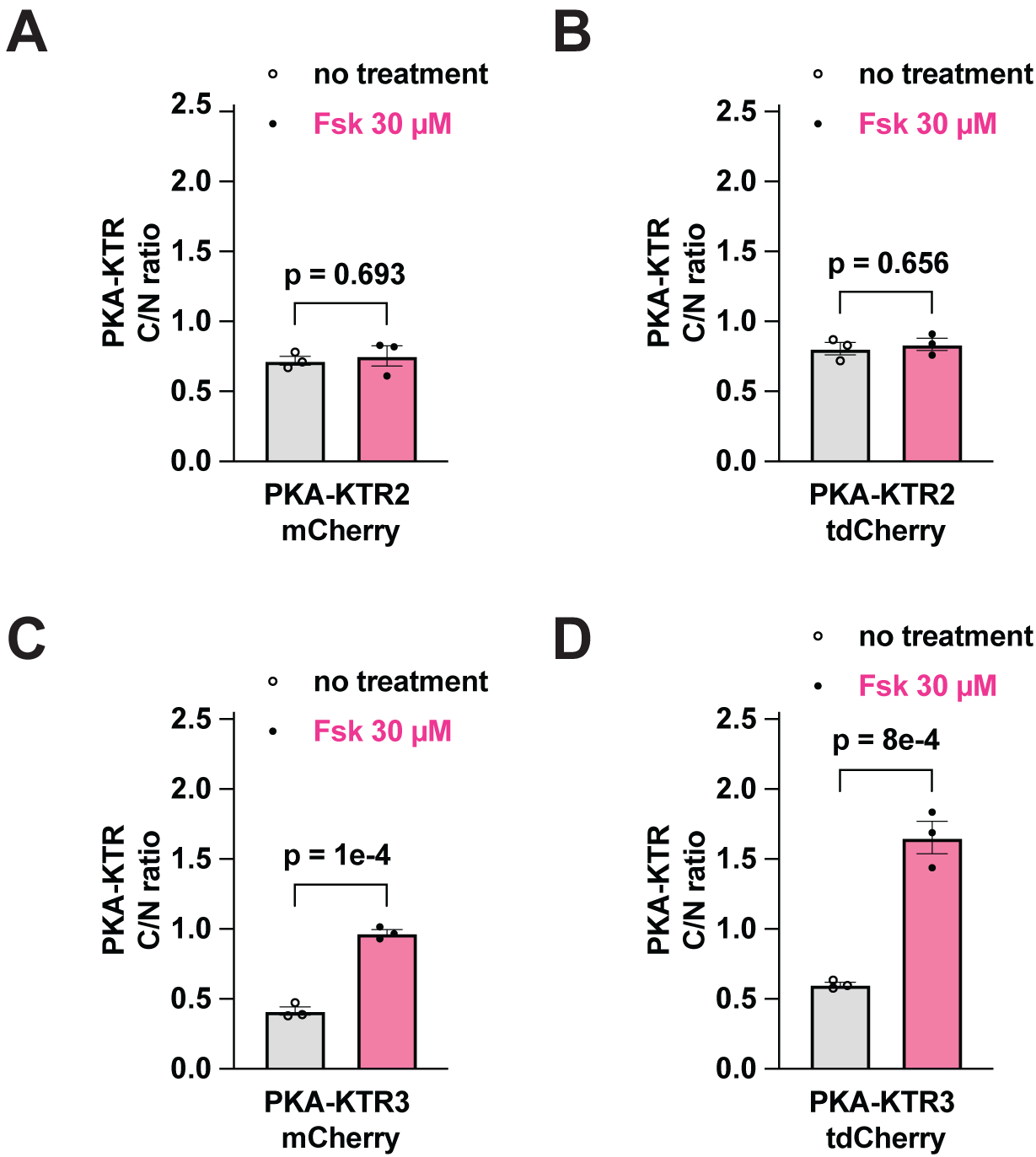
Increasing KTR size triples the dynamic range of PKA-KTR3. HEK293 cells were transfected with plasmid vectors designed to express fusion proteins comprised of the sensor domains of PKA-KTR2(*28*) or PKA-KTR3(*29*) appended to the N-terminus of either mCherry, tdCherry. Two days later, cells were imaged by fluorescence microscopy prior to or after a 30 min. incubation in 30 mM Fsk, followed by calculation of C/N ratios in 10 cells in independent regions of interest (ROIs) from three biological replicates, shown here in bar graph form. (**A, B**) Both PKA-KTR2/mCherry, a ∼34 kDa proteins, and PKA-KTR2/tdCherry, a ∼64 kDa protein, had resting C/N ratios <1, and neither showed a Fsk-induced translocation to the cytoplasm. (**C**) PKA-KTR3/mCherry, a ∼34 kDa protein, displayed a low resting C/N ratio and moved to the cytoplasm in response to Fsk, confirming that the PKA-KTR3 sensor domain is a useful reporter of PKA activity. However, Fsk-induced C/N ratio of PKA-KTR3/mCherry remained <1, and its dynamic range was relatively narrow, only ∼2-fold. (**D**) In contrast, we found that PKA-KTR3/tdCherry, ∼64 kDa, had a broader dynamic range, ∼3-fold, showing once again that increasing KTR size can be sufficient to improve KTR performance characteristics. Data are from a minimum of three independent biological replicates. ANOVA *p* values are denoted by *** <0.001.

**Figure S5.**
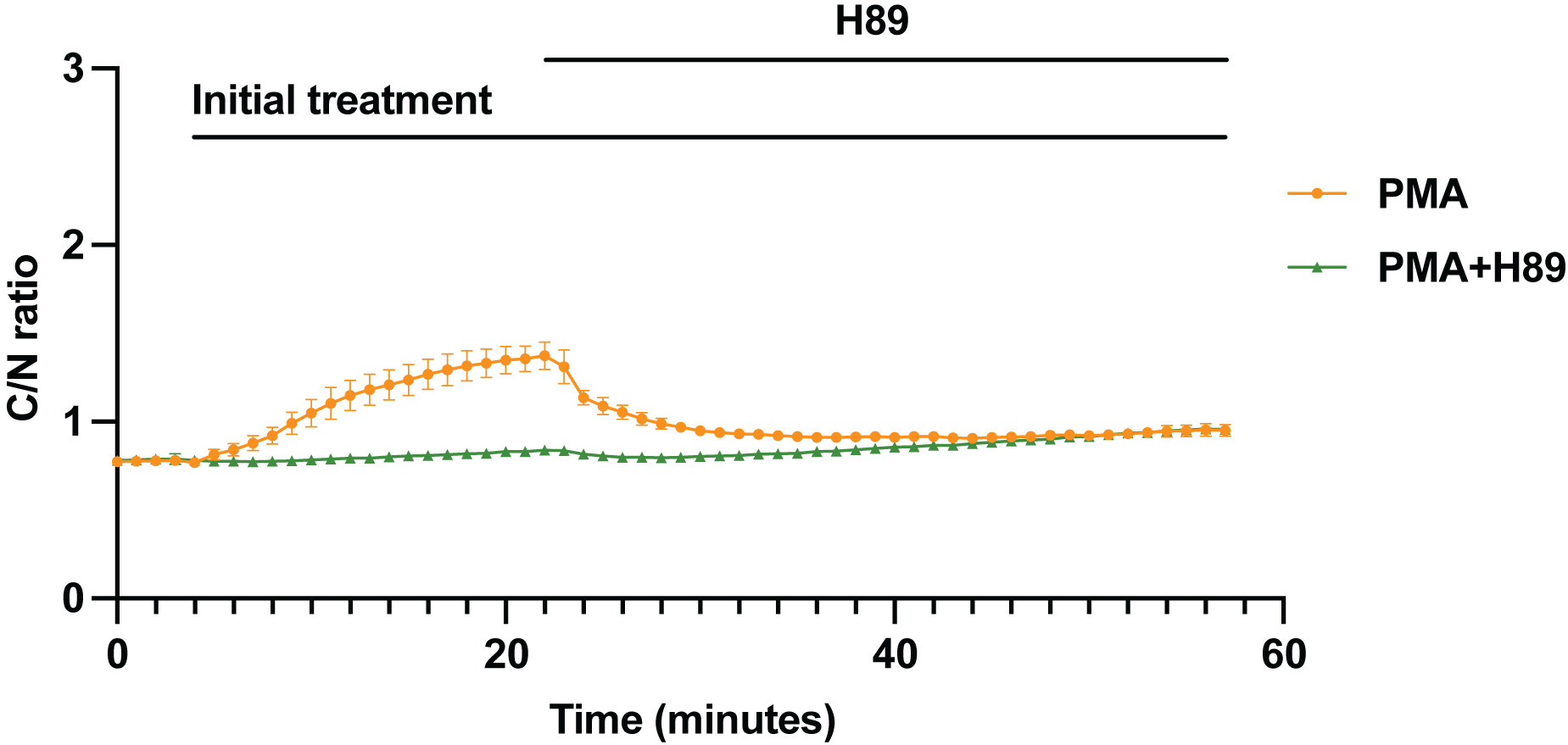
PMA induces a mild activation of PKA. Plot of C/N ratios for ePKA-KTR1.2/tdTomato C/N at every minute in ePKA-KTR1.2/tdTomato-expressing HEK293 cells exposed to (orange) PMA at t = 3 min, followed by the addition of the PKA inhibitor H89 at t = 23, or (green) PMA and H89 at t = 3, followed by addition of H89 again at t = 23 min.

**Table S1.**
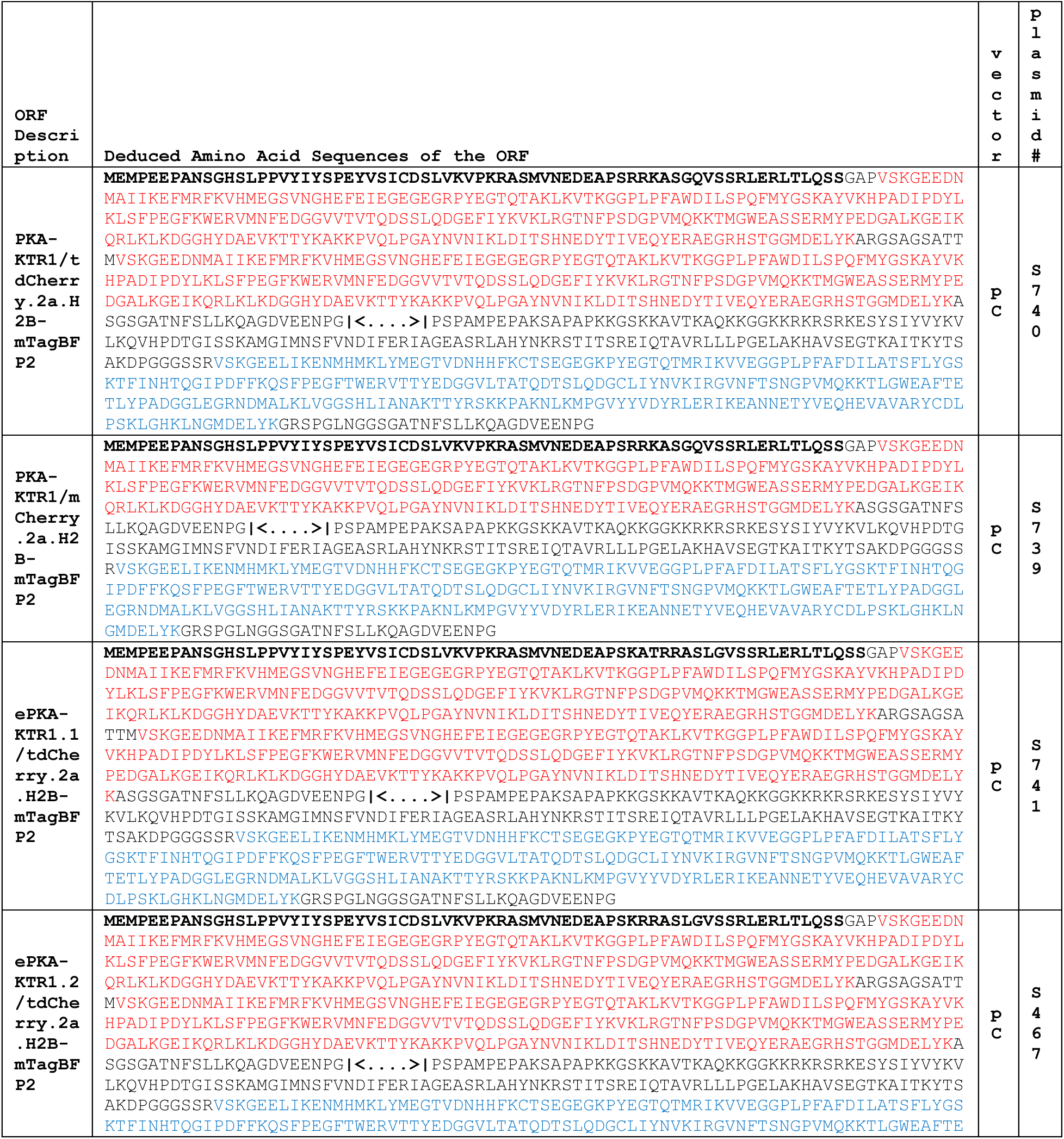
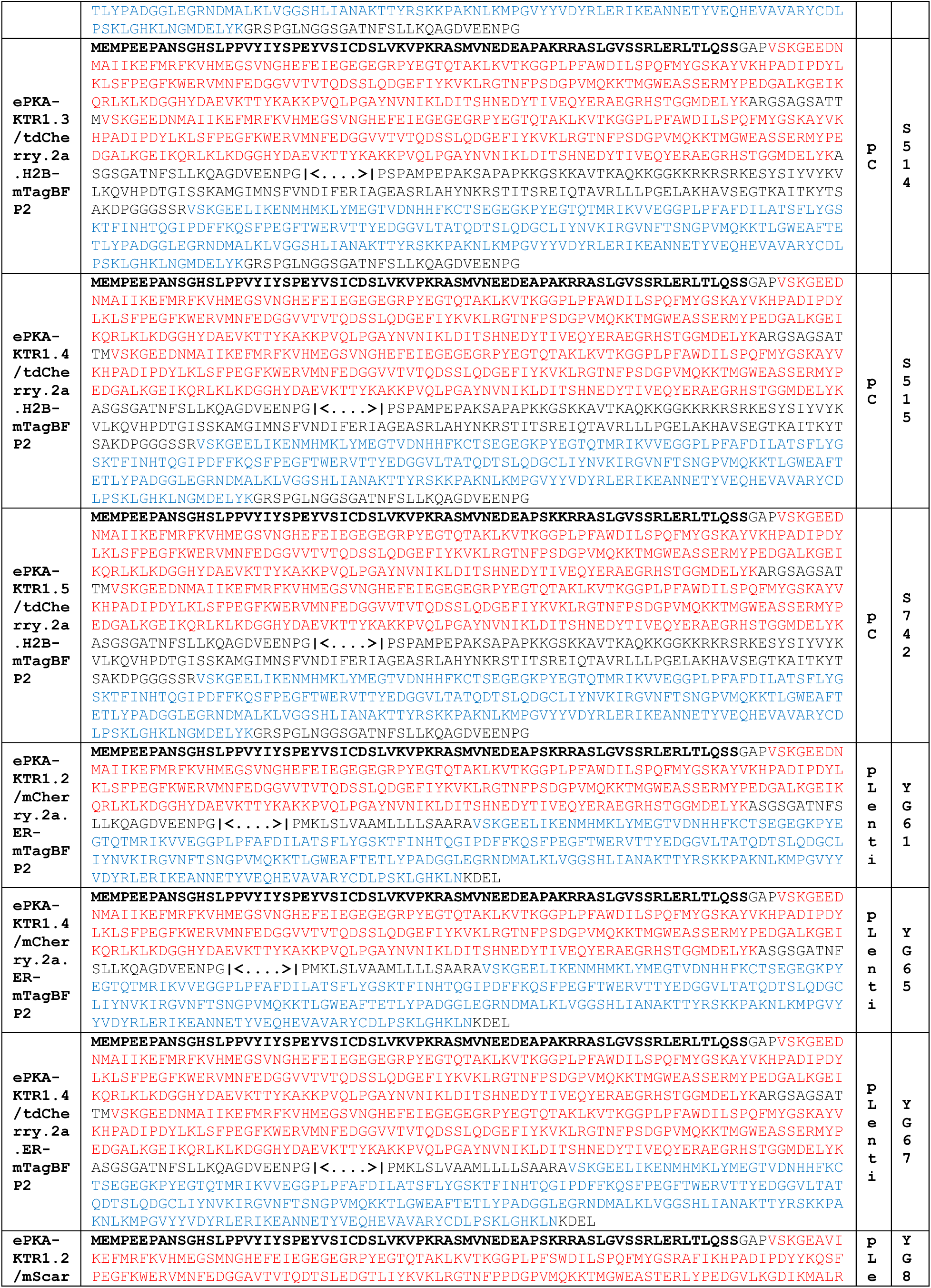
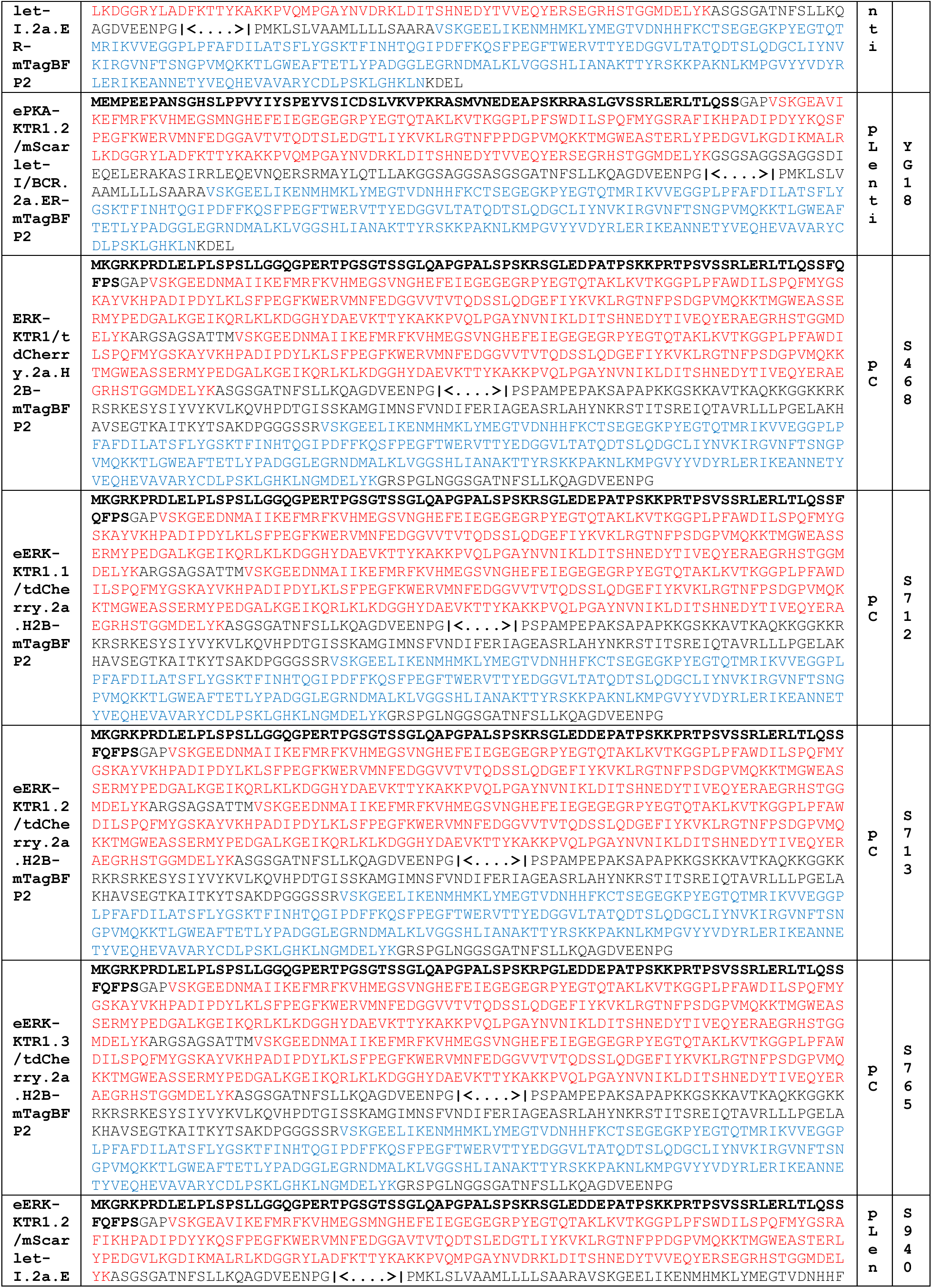
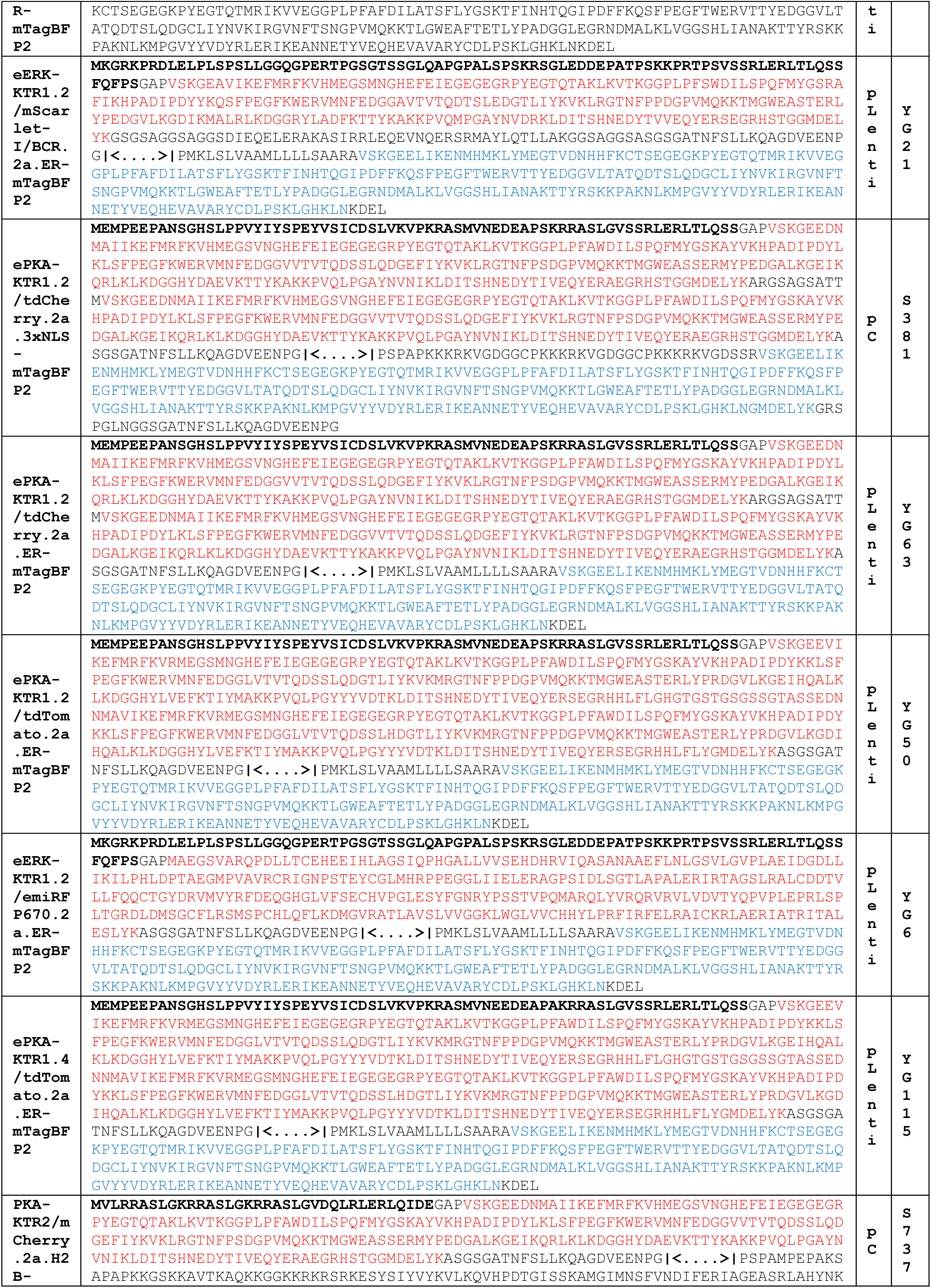
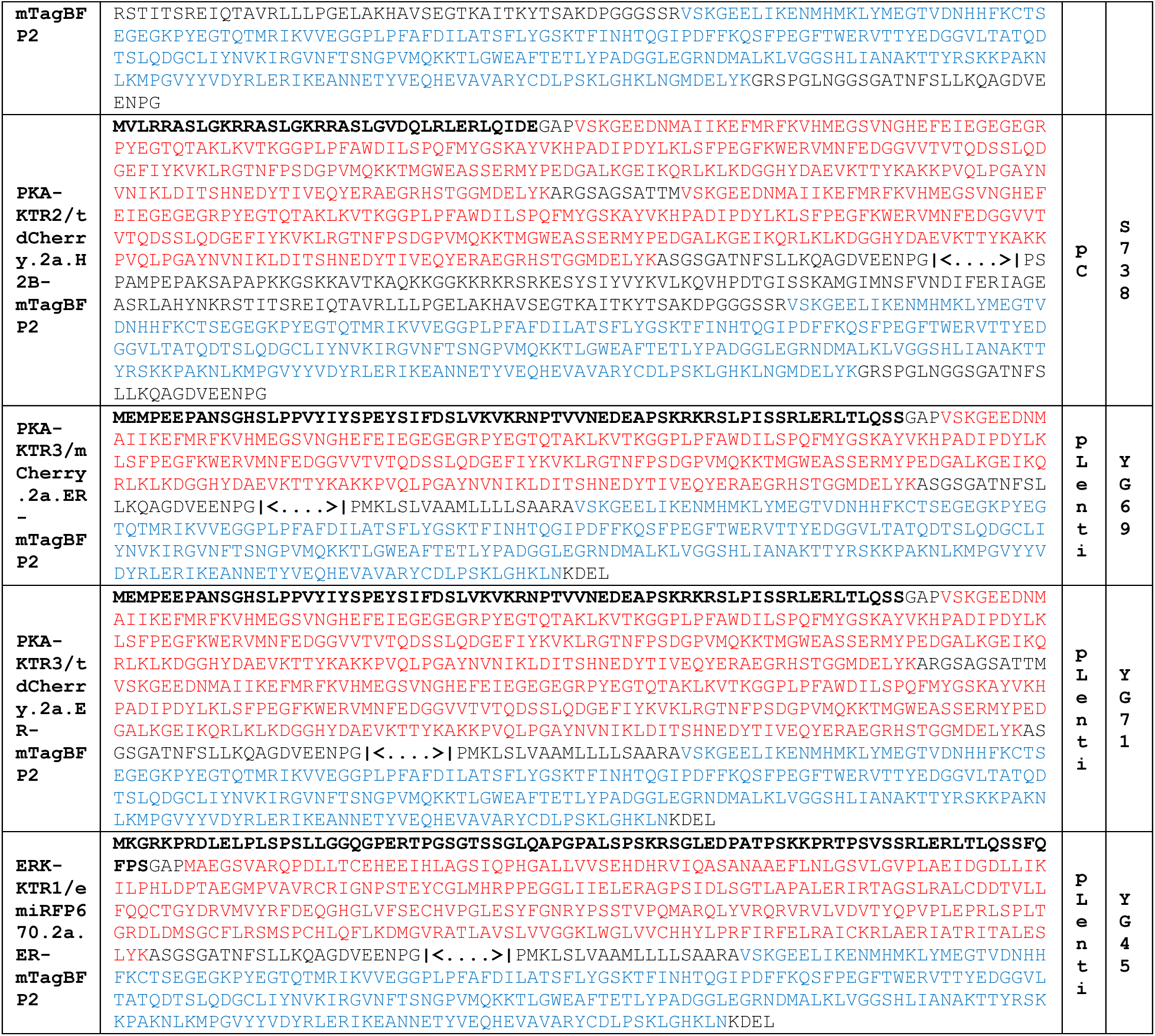
Description, amino acid sequences, vector type, and plasmid number. Vectors pC and pLenti were described previously(*62*). Amino acid sequences are represented in single letter code, with bold, black lettering for kinase sensor domains, red lettering for red and infrared fluorescent proteins, blue lettering for mTagBFP2, and black, unbolded lettering for other sequence elements (i.e. linker sequences, histone H2B sequence, C-terminal extensions, signal sequence, ER retrieval signal, and 3xNLS). The notation **|<….>|** denotes the position in the viral 2a peptide where the ribosome fails to make the peptide bond (between the upstream glycine and the downstream proline), resulting in the release of the upstream red or infrared fluorescent protein and the continued translation of the downstream blue fluorescent protein.

**Table S2.** List of plasmid numbers and the ORF they express.

| **Name** | **Description (ORF expressed)** | **vector backbone** | **promoter** |
| --- | --- | --- | --- |
| S739 | PKA-KTR1-mCherry 2a H2B-mTagBFP2 | pC (plasmid) | CMV |
| S777 | mCherry-PKA-KTR2 2a H2B-mTagBFP2 | pC (plasmid) | CMV |
| S740 | PKA-KTR1-tdCherry 2a H2B-mTagBFP2 | pC (plasmid) | CMV |
| S738 | PKA-KTR2-tdCherry 2a H2B-mTagBFP2 | pC (plasmid) | CMV |
| S741 | ePKA-KTR1.1-tdCherry 2a H2B-mTagBFP2 | pC (plasmid) | CMV |
| S467 | ePKA-KTR1.2-tdCherry 2a H2B-mTagBFP2 | pC (plasmid) | CMV |
| S514 | ePKA-KTR1.3-tdCherry 2a H2B-mTagBFP2 | pC (plasmid) | CMV |
| S515 | ePKA-KTR1.4-tdCherry 2a H2B-mTagBFP2 | pC (plasmid) | CMV |
| S742 | ePKA-KTR1.5-tdCherry 2a H2B-mTagBFP2 | pC (plasmid) | CMV |
| YG63 | ePKA-KTR1.2-tdCherry 2a SS-mTagBFP2-KDEL | pLenti (lentiviral vector) | CMV |
| S381 | ePKA-KTR1.2-tdCherry 2a 3xNLS-mTagBFP2 | pC (plasmid) | CMV |
| YG61 | ePKA-KTR1.2-mCherry 2a SS-mTagBFP2-KDEL | pLenti (lentiviral vector) | CMV |
| YG65 | ePKA-KTR1.4-mCherry 2a SS-mTagBFP2-KDEL | Lentiviral | CMV |
| YG67 | ePKA-KTR1.4-tdCherry 2a SS-mTagBFP2-KDEL | Lentiviral | CMV |
| YG8 | ePKA-KTR1.2-mScarlet 2a SS-mTagBFP2-KDEL | Lentiviral | CMV |
| YG18 | ePKA-KTR1.2-mScarlet-BCR 2a SS-mTagBFP2-KDEL | Lentiviral | CMV |
| S468 | ERK-KTR1-tdCherry 2a H2B-mTagBFP2 | pC (plasmid) | CMV |
| S712 | eERK-KTR1.1-tdCherry 2a H2B-mTagBFP2 | pC (plasmid) | CMV |
| S713 | eERK-KTR1.2-tdCherry 2a H2B-mTagBFP2 | pC (plasmid) | CMV |
| S765 | eERK-KTR1.3-tdCherry 2a H2B-mTagBFP2 | pC (plasmid) | CMV |
| S940 | eERK-KTR1.2-mScarlet 2a SS-mTagBFP2-KDEL | Lentiviral | CMV |
| YG21 | eERK-KTR1.2-mScarlet-BCR 2a SS-mTagBFP2-KDEL | Lentiviral | CMV |
| YG6 | eERK-KTR1.2-emiRFP670 2a SS-mTagBFP2-KDEL | pLenti (lentiviral vector) | CMV |
| YG45 | ERK-KTR1-emiRFP670 2a SS-mTagBFP2-KDEL | pLenti (lentiviral vector) | CMV |
| YG50 | ePKA-KTR1.2-tdTomato 2a SS-mTagBFP2-KDEL | pLenti (lentiviral vector) | CMV |
| YG114 | PKA-KTR3-tdTomato 2a SS-mTagBFP2-KDEL | pLenti (lentiviral vector) | CMV |
| YG115 | ePKA-KTR1.4-tdTomato 2a SS-mTagBFP2-KDEL | pLenti (lentiviral vector) | CMV |
| S1137 | jGCaMP8s | Lentiviral | CMV |
| YG160 | eERK-KTR1.2 (mutated)-emiRFP670 2a SS-mTagBFP2-KDEL | pLenti (lentiviral vector) | CMV |

## Structured Methods - Reagents and Tools Table

*Instructions: Please complete the relevant fields below, adding rows as needed. The following page provides an example of a completed table and additional instruction for entering your data in the table*.

| Reagent/Resource | Reference or Source | Identifier or Catalog Number |
| --- | --- | --- |
| **Experimental Models** |  |  |
| HEK-293 cells (H. sapiens) | ATCC | CRL-1573 |
| Dorsal root ganglion neurons | C57BL/6 (Jackson Lab) | 000664 |
| **Recombinant DNA** |  |  |
| psPAX2 | Addgene | 12260 |
| VSV.G | Addgene | 14888 |
| PKA-KTR1/tdCherry.2a.H2B-mTagBFP2 | This study | N/A |
| PKA-KTR1/mCherry.2a.H2B-mTagBFP2 | This study | N/A |
| ePKA-KTR1.1/tdCherry.2a.H2B-mTagBFP2 | This study | N/A |
| ePKA-KTR1.2/tdCherry.2a.H2B-mTagBFP2 | This study | N/A |
| ePKA-KTR1.3/tdCherry.2a.H2B-mTagBFP2 | This study | N/A |
| ePKA-KTR1.4/tdCherry.2a.H2B-mTagBFP2 | This study | N/A |
| ePKA-KTR1.5/tdCherry.2a.H2B-mTagBFP2 | This study | N/A |
| ePKA-KTR1.2/mCherry.2a.ER-mTagBFP2 | This study | N/A |
| ePKA-KTR1.4/mCherry.2a.ER-mTagBFP2 | This study | N/A |
| ePKA-KTR1.4/tdCherry.2a.ER-mTagBFP2 | This study | N/A |
| ePKA-KTR1.2/mScarlet-I.2a.ER-mTagBFP2 | This study | N/A |
| ePKA-KTR1.2/mScarlet-I/BCR.2a.ER-mTagBFP2 | This study | N/A |
| ERK-KTR1/tdCherry.2a.H2B-mTagBFP2 | This study | N/A |
| eERK-KTR1.1/tdCherry.2a.H2B-mTagBFP2 | This study | N/A |
| eERK-KTR1.2/tdCherry.2a.H2B-mTagBFP2 | This study | N/A |
| eERK-KTR1.3/tdCherry.2a.H2B-mTagBFP2 | This study | N/A |
| eERK-KTR1.2/mScarlet-I.2a.ER-mTagBFP2 | This study | N/A |
| eERK-KTR1.2/mScarlet-I/BCR.2a.ER-mTagBFP2 | This study | N/A |
| ePKA-KTR1.2/tdCherry.2a.3xNLS-mTagBFP2 | This study | N/A |
| ePKA-KTR1.2/tdCherry.2a.ER-mTagBFP2 | This study | N/A |
| ePKA-KTR1.2/tdTomato.2a.ER-mTagBFP2 | This study | N/A |
| eERK-KTR1.2/emiRFP670.2a.ER-mTagBFP2 | This study | N/A |
| ePKA-KTR1.4/tdTomato.2a.ER-mTagBFP2 | This study | N/A |
| PKA-KTR2/mCherry.2a.H2B-mTagBFP2 | This study | N/A |
| PKA-KTR2/tdCherry.2a.H2B-mTagBFP2 | This study | N/A |
| PKA-KTR3/mCherry.2a.ER-mTagBFP2 | This study | N/A |
| PKA-KTR3/tdCherry.2a.ER-mTagBFP2 | This study | N/A |
| ERK-KTR1/emiRFP670.2a.ER-mTagBFP2 | This study | N/A |
| Additional plasmids and more information | This study | Table S1 |
| **Antibodies** |  |  |
| Rabbit anti-phospho-ERK1/2 | Cell Signaling Technology | 9101S |
| Rabbit anti-phospho-PKA R2 (Ser99) | Abcam | ab32390 |
| Goat anti-rabbit Alexa488 | Jackson ImmunoResearch | AB_2338046 |
| **Oligonucleotides and other sequence-based reagents** |  |  |
| Geneblocks and primers | Integrated DNA Technologies | Table S1 |
| **Chemicals, Enzymes and other reagents** |  |  |
| Phusion High-Fidelity DNA Polymerase | ThermoFisher | F530L |
| T4 DNA Ligase (5 U/μL) | ThermoFisher | EL0011 |
| Petri Dishes | Falcon | 351029 |
| DMEM, high glucose | ThermoFisher | 11965118 |
| Trypsin-EDTA (0.25%), phenol red | ThermoFisher | 25200114 |
| Fetal Bovine Serum | ThermoFisher | 16000044 |
| 24-well cell culture plates | Denville | 1156F00 |
| TC-treated Culture Dish (100 mm) | Corning | 430167 |
| Hanks' Balanced Salt Solution (HBSS) | ThermoFisher | 14175103 |
| 96 Well Black Plate, Coverglass Bottom, CC2 Surface | ThermoFisher | 160376 |
| μ-Slide 8 Well high Glass Bottom | Ibidi | 80807 |
| μ-Slide 18 Well Glass Bottom | Ibidi | 81817 |
| A23187 | Sigma-Aldrich | C7522 |
| ATP | Sigma-Aldrich | A26209 |
| Forskolin | Cayman Chemical | 11018 |
| H89 | MedChemExpress | HY-15979A |
| EGF | Gibco | PHG0311L |
| VX-11e | SelleckChem | S7709 |
| leptomycin B | Sigma-Aldrich | L2913 |
| Phorbol 12-myristate 13-acetate | Sigma-Aldrich | P1585 |
| Neurobasal™ Plus Medium | Gibco | A3582901 |
| B-27™ Plus Supplement | Gibco | A3582801 |
| Poly-L-lysine solution | Sigma-Aldrich | P8920-100ML |
| Laminin | Sigma-Aldrich | L2020 |
| Papain, suspension | Worthington | LS003126 |
| Bovine Serum Albumin | Sigma-Aldrich | A-9418 |
| TM Liberase | Roche | 5401119001 |
| TL Liberase | Roche | 5401020001 |
| Pen/Strep (100X) | ThermoFisher | 15140122 |
| Lipofectamine™ 3000 | ThermoFisher | L3000015 |
| Open-Top Thinwall Ultra-Clear Tube | Beckman-Coulter | C14292 |
| 50 mL Luer-Lok™ Syringe | BD | 309653 |
| Syringe Filter (0.45 µm) | Millipore | SLHVR33RS |
| 8-mm cover glasses | Electron Microscopy Sciences | 7229608 |
| QIAprep Spin Miniprep Kit | Qiagen | 27106 |
| Wizard® Plus Midipreps DNA Purification System | Promega | A7640 |
| Ampicillin sodium salt | Sigma-Aldrich | A8351 |
| **Software** |  |  |
| GraphPad Prism 10 | https://www.graphpad.com/ |  |
| SnapGene 7 | https://www.snapgene.com/ |  |
| ImageJ (Fiji) | https://fiji.sc/ |  |
| NIS Elements Advanced Research v4.60 | https://www.microscope.healthcare.nikon.com/products/software/nis-elements |  |
| EVOS M7000 Cell Imaging System Software | https://www.thermofisher.com/us/en/home/technical-resources/software-downloads/evos-m7000-imaging-system-software-download.html |  |
| **Other** |  |  |
| EVOS M7000 Imaging System | ThermoFisher | AMF7000 |
| EVOS Onstage Incubator | ThermoFisher | AMC2000 |
| Optima L-90K ultracentrifuge | Beckman-Coulter |  |
| SW-32 Ti rotor | Beckman-Coulter |  |
| Sony MA900 Cell Sorter | Sony |  |

